# Ribosome abundance, not paralogue composition, is essential for germline development

**DOI:** 10.1101/2024.11.14.623487

**Authors:** Katarina Z A Grobicki, Daniel Gebert, Carol Sun, Felipe Karam Teixeira

**Affiliations:** Department of Genetics, University of Cambridge, Downing Street, CB2 3EH, Cambridge, UK; Department of Physiology, Development, and Neuroscience, University of Cambridge, Downing Street, CB2 3DY, Cambridge, UK

## Abstract

Ribosomes catalyse all protein synthesis, and mutations altering their levels and function underlie many developmental diseases and cancer. Historically considered to be invariant machines, ribosomes differ in composition between tissues and developmental stages, incorporating a diversity of ribosomal proteins (RP) encoded by duplicated paralogous genes. Here, we use *Drosophila* to systematically investigate the origins and functions of non-canonical RP paralogues. We show that new paralogues mainly originated through retroposition, and that only a few new copies retain coding capacity over time. Although transcriptionally active non-canonical RP paralogues often present tissue-specific expression, we show that the majority of those are not required for either viability or fertility in *Drosophila melanogaster*. The only exception, RpS5b, which is required for oogenesis, is functionally interchangeable with its canonical paralogue, indicating that the *RpS5b^-/-^* phenotype results from insufficient ribosomes rather than the absence of an RpS5b-specific, functionally-specialised ribosome. Altogether our results provide evidence that, instead of new functions, RP gene duplications provide a means to regulate ribosome levels during development.

## Introduction

Ribosomes are RNA-protein machines responsible for protein translation across all domains of life. As such, production of sufficient functional ribosomes is essential for growth and development. In humans, mutations affecting genes encoding ribosomal proteins (RPs) or ribosome biogenesis factors lead to a group of rare developmental diseases known as ribosomopathies ^1–3^. Despite arising from defects affecting such a ubiquitous complex, ribosomopathies produce remarkably tissue-specific phenotypes. For example, mutations in *treacle*, required for rRNA transcription and 18S methylation, primarily affect neural crest cells to cause Treacher-Collins syndrome (Teng *et al*., 2013; Valdez *et al*., 2004), while mutations in a number of RPs primarily affect haematopoiesis and cause Diamond-Blackfan anaemia ^2,4,5^ and mutations in *snoRNA U8* exclusively affect the white matter of the brain ^6,7^. In contrast to the progress made in building a detailed biochemical and structural model for the process of translation, our understanding of how the activity of ribosomes and translation impacts organismal biology remains superficial.

One appealing avenue to explain how ribosomopathies can affect different cell types is the fact that ribosomes, long-thought to be invariant machines, are actually more heterogeneous than previously appreciated ^8^. This heterogeneity can arise through differences in rRNA modifications, post-translational modifications on RPs, assembly of RP paralogues, or the association of ribosomes with different accessory factors ^9^. Differences in ribosome composition have been hypothesised to determine the speed or fidelity of translation, the sub-cellular localisation of ribosomes, or even a preference for translating specific subsets of mRNA transcripts ^9^. At an organism level, mounting evidence indicates that ribosome heterogeneity exists between cell types and at different developmental stages ^10–13^, and that these differences may influence cellular function. For instance, a combination of mouse genetics and cryo-electron microscopy have demonstrated that a testes-specific RpL39 paralogue, RpL39-like, is required for the co-translational folding of key spermatogenesis proteins ^14,15^. Despite some progress, the evidence that ribosome heterogeneity has a direct functional effect during development is mostly anecdotal, and this hypothesis has not been tested systematically.

The use of *Drosophila melanogaster* as a model organism for over 100 years has resulted in detailed characterisation of development, a well-annotated genome, and an arsenal of genetic tools, making it an ideal organism to genetically investigate ribosome heterogeneity. RP mutations in *Drosophila* were first identified due to their characteristic *minute* phenotype, comprising delayed development, short thin bristles, and reduced fertility and viability (Bridges and Morgan, 1923). Continued research has shown that precise regulation of ribosome biogenesis and function is essential for *Drosophila* development. For example, in both neuroblasts ^17^ and germline stem cells ^18,19^, stem cells maintain higher levels of ribosome biogenesis but lower translation compared to their daughter cells, while the inverse is required for differentiation ^19,20^. Ribosome heterogeneity is also being increasingly appreciated in *Drosophila*, with rRNA 2′-O-methylation ^21^ and pseudouridylation ^22^ modifications recently being profiled in a number of tissues. A number of RP paralogues and translation initiation factors were shown to have enriched expression in germline stem cells ^23^. These paralogues have since been found to be assembled into translating ribosomes in the ovary and the testis ^24^, raising the intriguing possibility of germline-specific ribosomes playing a key role in gametogenesis and embryo development.

Here, we used the genetically-tractable model *Drosophila* to systematically examine RP paralogues with respect to both evolution and function. We show that RP genes are frequently duplicated, however, only in a few cases are the resulting duplicates retained or expressed. We individually mutated 11 RP paralogues that are expressed in *D. melanogaster*, and found that 10 were not required for viability or fertility, producing no discernible phenotype under laboratory conditions. Only one RP paralogue, *RpS5b*, was required for female fertility, with mutation leading to ectopic TORC1 activity, metabolic remodelling and germ cell death. However, we reveal that the same phenotype can be induced by depleting germline ribosomes by alternative methods, and demonstrate that RpS5b and its paralogue RpS5a are functionally interchangeable. Together, these results show that the ribosome heterogeneity provided by RP paralogues does not generally impart functional differences and is not required for development. Instead, we demonstrate that RP paralogues can provide a key role in sustaining ribosome abundance in specific cellular contexts, and with that becoming essential for normal development.

## Results

### RP genes are frequently duplicated, mostly by retroposition

Due to their essentiality, RP genes are under strict purifying selection ^25^; however, duplications of RP genes have the potential to allow the emergence of subfunctionalisation or neofunctionalisation. Although no systematic analysis of RP gene duplication has been carried out to date, expression evidence has been used to identify 14 RP gene duplications in the *D. melanogaster* genome ^26,27^. To systematically identify all the copies of RP genes in a given genome, we designed a BLAST-based pipeline and analysed the genomes of 12 *Drosophila* species, representing approximately 73 million years (My) of evolution. First, amino acid sequences of 79 “canonical” RPs from *Drosophila melanogaster* were used as queries to perform tBLASTn searches. We relied on the fact that, except for individual gene duplications, genetic material is rarely exchanged between chromosome arms in the *Drosophila* genus ^28–30^ to reconstruct the history of the duplication events and place them within the phylogeny. This was achieved by determining the genomic location of each identified loci in the highly-contiguous genomes that were assembled using long-read sequencing and chromatin conformation data ^31^, the loci structure of each identified copy, and the sequence similarity scores between copies (see Methods). The minimum age of a given duplication event was estimated based on the most parsimonious phylogenetic explanation ^32^. Finally, we performed loci structure comparisons between new copies and their “parent gene” to classify each duplication event as either a retroposition (loss of parental introns), a DNA-mediated duplication (retention of intron-exon structure), or else unknown (in cases where the “parent gene” is intronless).

In total, we identified 1373 RP loci in the genomes of the 12 *Drosophila* species studied, corresponding to the 79 “canonical” genes and 388 independent duplication events (Figure 1A). Given that most of the 12 species analysed here diverged >20My ago, it is not surprising that 83% of the identified duplication events were shown to be species-specific (Figure 1B). However, and despite the fact that *D. melanogaster* protein sequences were used as bait, we did not observe any correlation between the divergence time from *D. melanogaster* and the number of duplicates identified (R^2^=0.048; Figure S1A). We were able to classify the mechanism of duplication for about 2/3 of the events, with 87% of these (N=234) being a result of retroposition and the remaining 13% (N=35) occurring by DNA-mediated duplication. The estimated minimum age of duplication events differed between the two modes of duplications, with DNA-mediated duplication events being older on average than retroduplications (Figure 1C).

**Figure 1.**
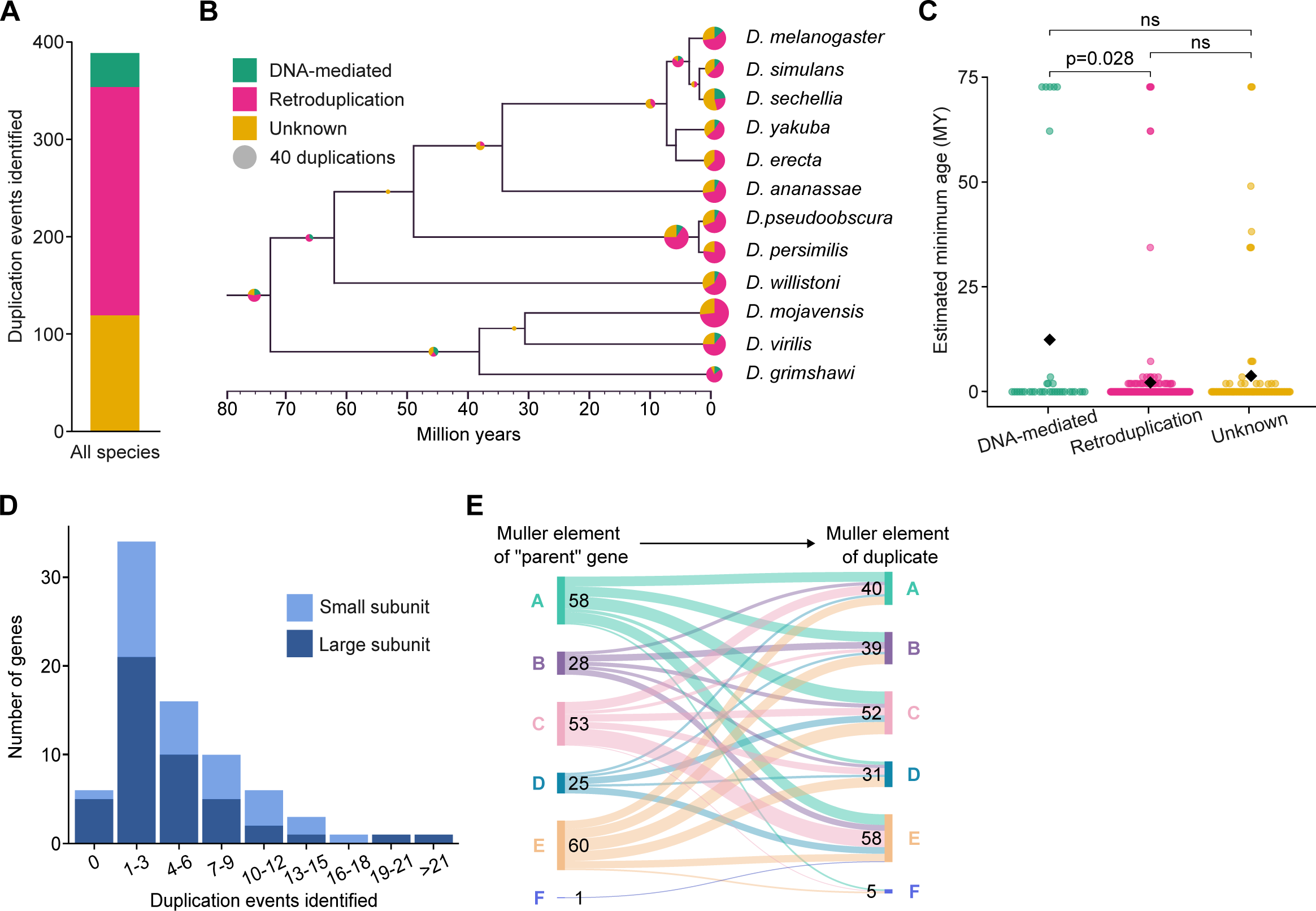
The majority of RP gene duplications occur by retroposition and are evolutionarily recent. (A) All duplication events identified in the genomes of 12 *Drosophila* species, coloured by mode of duplication as indicated in B. (**B**) All duplication events visualised on phylogenetic tree (scale based on Thomas and Hahn, 2017). Size of circle represents number of duplication events; colour represents mode of duplication. (**C**) Estimated minimum age of duplication events based on phylogeny. Black diamonds represent means; *P*-values from Mann Whitney U test (Wilcoxon rank sum test) are indicated at the top. (**D**) Histogram of number of duplication events identified for each RP gene. (**E**) Relationships between the Muller element of “parent” genes and retroposition-derived duplicates. N=225 (all retroposition events where the Muller element of both the “parent” gene and the duplicate could be assigned; 96% of identified retroposition events).

We also assessed whether there were biases for which “canonical” RP genes are duplicated. The number of duplication events linked to any given “canonical” RP gene varied greatly, with most genes being associated with 1-3 duplication events, while one was associated with as many as 24 events (Figure 1D). We did not observe any significant difference between the number of duplications associated with RP genes encoding either large or small ribosomal subunit proteins, or based on their position within the ribosome structure (Figure S1B). However, the total number of duplication events associated with each gene showed a moderate correlation with the mRNA expression level of the “canonical” RP genes in *D. melanogaster* ovary and testis (Figure S1C-D). Altogether, these results indicate that most duplications we identified are caused by retroposition events and suggest that mRNA concentration in the germline is an important factor influencing the frequency of events.

Finally, we examined the relationship between the genome distribution of the retroposition-mediated duplicated RP loci in relation to the location of the “parent gene” (N=225 retroposition events). This analysis did not reveal any specific trend, with the patterns of retroduplication not differing significantly from the expected random distribution (Figure 1E). Previous analysis of *Drosophila* retroduplicated genes identified a trend whereby a large proportion of retroduplicates found on autosomes have originated from “parent genes” located on the X-chromosome, and the new autosomal copies showed an enriched expression in the testis ^33–36^. This led to the “out-of-X” hypothesis, proposing that new copies could “escape” X-chromosome inactivation during spermatogenesis and potentially evolve new functions in the male reproductive system. While these previous analyses focused exclusively on retroduplicates retaining transcriptional activity, our analysis encompasses all retroduplicates regardless of their potential for transcriptional activity. Together, these analyses support a scenario in which the “out-of-X” phenomenon arises from biases at the level of selection, rather than in the initial duplication events.

### In *D. melanogaster*, most RP paralogues are dispensable for viability or fertility

Our bioinformatic analysis identified 133 RP loci in the *D. melanogaster* genome (Figure 2A). 93 of those, including the 79 “canonical” RP genes, were previously annotated and showed to have transcriptional activity in at least one tissue ^26,27,37–39^. The manual analysis of the remaining 40 loci revealed that these were either fragmented, or the coding sequence contained premature codons, or showed little sequence conservation. Moreover, none of these loci showed transcriptional activity based on the analyses of the publicly available large datasets containing tissue-specific RNA-seq data ^26,37–39^, suggesting that they are likely pseudogenes.

**Figure 2.**
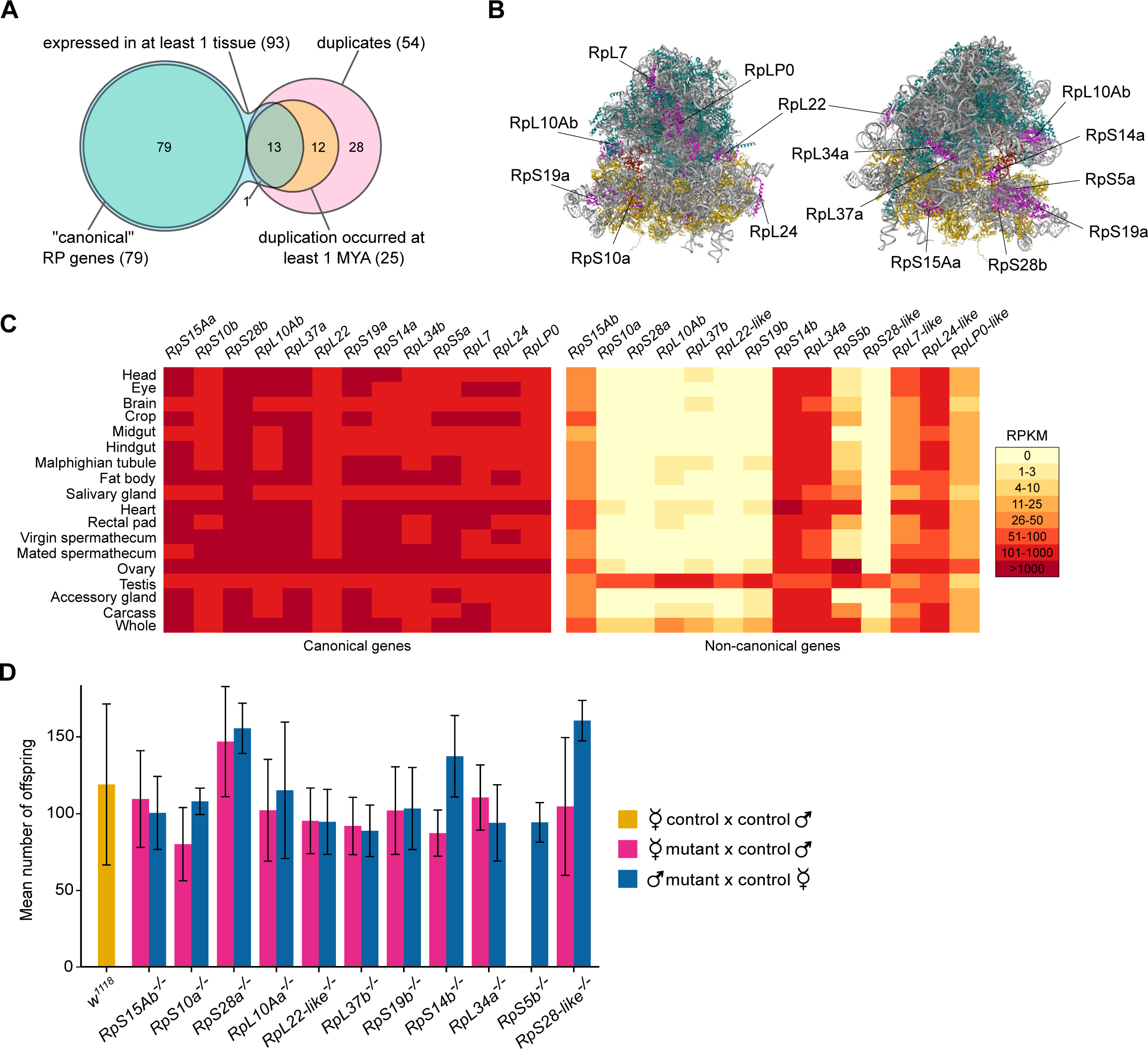
“Non-canonical” RP genes frequently show germline-specific expression, but the majority are not required for viability, germline development, or fertility in *D. melanogaster*. (**A**) Summary of distribution of the 133 RP loci identified in the *D. melanogaster* genome. (**B**) Structure of the *D. melanogaster* ribosome (Anger *et al*., 2013) depicting the locations of the 13 RPs encoded by two or more transcriptionally active paralogues (magenta). rRNA is grey, small subunit RPs are yellow, large subunit RPs are cyan. (**C**) Heat-map showing the mRNA accumulation levels (expressed as reads per kilobase per million mapped reads [RPKM] from publicly available RNA-seq data; Leader *et al*., 2018) of “canonical” and “non-canonical” RP genes in adult *D. melanogaster* tissues. (**D**) Number of F1 offspring eclosing from reciprocal crosses between homozygous “non-canonical” RP mutants and control (*w^1118^*) flies (pink bars indicate crosses between homozygous mutant females and control males; blue bars indicate crosses between control females and homozygous mutant males). Two different mutant allelic combinations were used for each “non-canonical” RP gene tested, with three independent replicates per allelic combination; six independent replicates were performed for the *w^1118^* x *w^1118^* control crosses (yellow bars). Error bars show standard deviation.

For 66 RPs, only one transcriptionally active gene was present in the *D. melanogaster* genome. For the remaining 13 RPs, we observed the presence of two, and in one case three, transcriptionally active paralogous genes (Figure 2B and S2A). Within each of these paralogue groups, at least one of the genes is expressed ubiquitously and at high levels (Figure 2C ^27,37^), and in most cases, homozygous loss-of-function mutations affecting these genes were previously shown to cause lethality (these genes are referred to henceforth as the “canonical” paralogues ^27^). For 9 of the 13 “canonical” RP genes, heterozygous loss-of-function mutations also cause the classic *minute* phenotype ^16,27^.

None of the other genes in the paralogue groups, referred to here as “non-canonical”, were previously assigned as *minute* genes. In contrast to the “canonical” paralogues, many “non-canonical” genes display tissue-specific expression patterns, particularly showing enriched expression in gonads (Figure 2C; ^23,26^). Two of the broadly-expressed “non-canonical” paralogues, *RpL24-like* and *RpLP0-like*, arose from ancient duplication events, and are conserved in organisms as distant as yeast and humans. Neither of these paralogues form part of the mature ribosome, but instead, they have evolved essential roles in ribosome biogenesis ^40,41^, so we did not consider them further in this study.

We used CRISPR/Cas9 mutagenesis to systematically generate 3-5 independent loss-of-function mutant alleles for 11 transcriptionally active “non-canonical” RP genes present in the *D. melanogaster* genome (gene and protein sequences of mutants in Table S1). Where available, paralogue-specific antibodies were used to validate knock-out mutants (Figure S2B-C). For all 11 genes, male and female homozygous loss-of-function mutants were viable to adulthood and did not display any features of the *minute* phenotype, demonstrating that these genes are not required for development or viability under laboratory conditions. Given that many of the “non-canonical” RP genes show enriched expression in the germline, we characterised the fertility of mutants by reciprocal crosses between homozygous mutants and control flies (*w^1118^*). The results revealed that male homozygous mutants were fertile for all 11 genes (Figure 2D), and in agreement, testes from all mutants appeared wild-type (Figure S3). Female homozygous mutants were fertile for 10 out of the 11 genes, however, *RpS5b^-/-^* females produced no offspring (Figure 2E). Phenotypic characterisation revealed that ovary morphology and germline development appeared wild type in all mutants, except for *RpS5b^-/-^* (Figure S3). Therefore, these results suggest that the majority of “non-canonical” RP genes are not required for viability, nor for male or female gonad development or function. *RpS5b* was the only exception, being required for female fertility.

### Germline *RpS5b* expression is required for oogenesis

*RpS5b* arose by DNA-mediated duplication at least 70 million years ago and is present in the genomes of 11 of the 12 *Drosophila* species examined here (gene and protein sequences in Table S2) - in *D. willistoni* the gene has been lost. In *D. virilis*, the *RpS5b* gene contains a premature stop codon and the mRNA is not detected in the ovary, however, in the 10 remaining species the *RpS5b* gene contains all the essential elements, and we confirmed the transcriptional activity of these in the ovaries of all the 8 species analysed ^31^.

In *D. melanogaster*, the RpS5a and RpS5b protein sequences share 80.4% sequence identity, with most differences occurring in the 40 amino acid N-terminal tail (Figure S4A). *RpS5a* is highly expressed in all tissues analysed, and homozygous mutations affecting *RpS5a* are lethal, whilst heterozygous mutants show a strong *minute* phenotype ^16,42^. *RpS5b* is highly expressed in the ovary and, to a lesser degree, the testes and heart ^37,39^. In ovaries, both RpS5a and RpS5b are expressed in the early germaria, where germline stem cell differentiation occurs (Figure S4B ^43^). However, from region 2b onwards, their expression patterns diverge, with RpS5a becoming preferentially expressed in somatic cells in comparison to the germline, while RpS5b is expressed almost exclusively in the germline (Figure S4B ^43^). Despite this divergence in expression pattern, Western blotting analyses indicated that both RpS5a and RpS5b are maternally deposited into eggs and therefore expressed during oogenesis (Figure S4C). Immunofluorescence of fixed embryos shows that maternally deposited RpS5a and RpS5b are broadly distributed in the syncytial embryo (Figure S4D).

As *RpS5b^-/-^* females were sterile, we used immunofluorescence to examine the ovary phenotype and observed that all mid-oogenesis egg chambers collapse in homozygous mutant ovaries and no mature eggs are produced (Figure 3A). Somatic follicle cells also showed abnormal organisation and multilayering, consistent with previous analyses using independent mutants ^43,44^. To examine whether apoptosis was activated, we stained *RpS5b^-/-^* ovaries with an antibody against cleaved Caspase-3 ^45^ and found strong signal in dying germ cells, but not somatic cells (Figure 3B). Finally, we determined whether RpS5b is required in both soma and germline by employing the FLP-FRT system ^46^ to produce *RpS5b^-/-^* clones in both cell lineages. As previously observed ^43^, our analysis demonstrated that *RpS5b* expression is only required in the germline (Figure S5A-B), and that the somatic multilayering phenotype is non-autonomously induced, likely resulting from disrupted germline-soma communication downstream of *RpS5b*.

**Figure 3.**
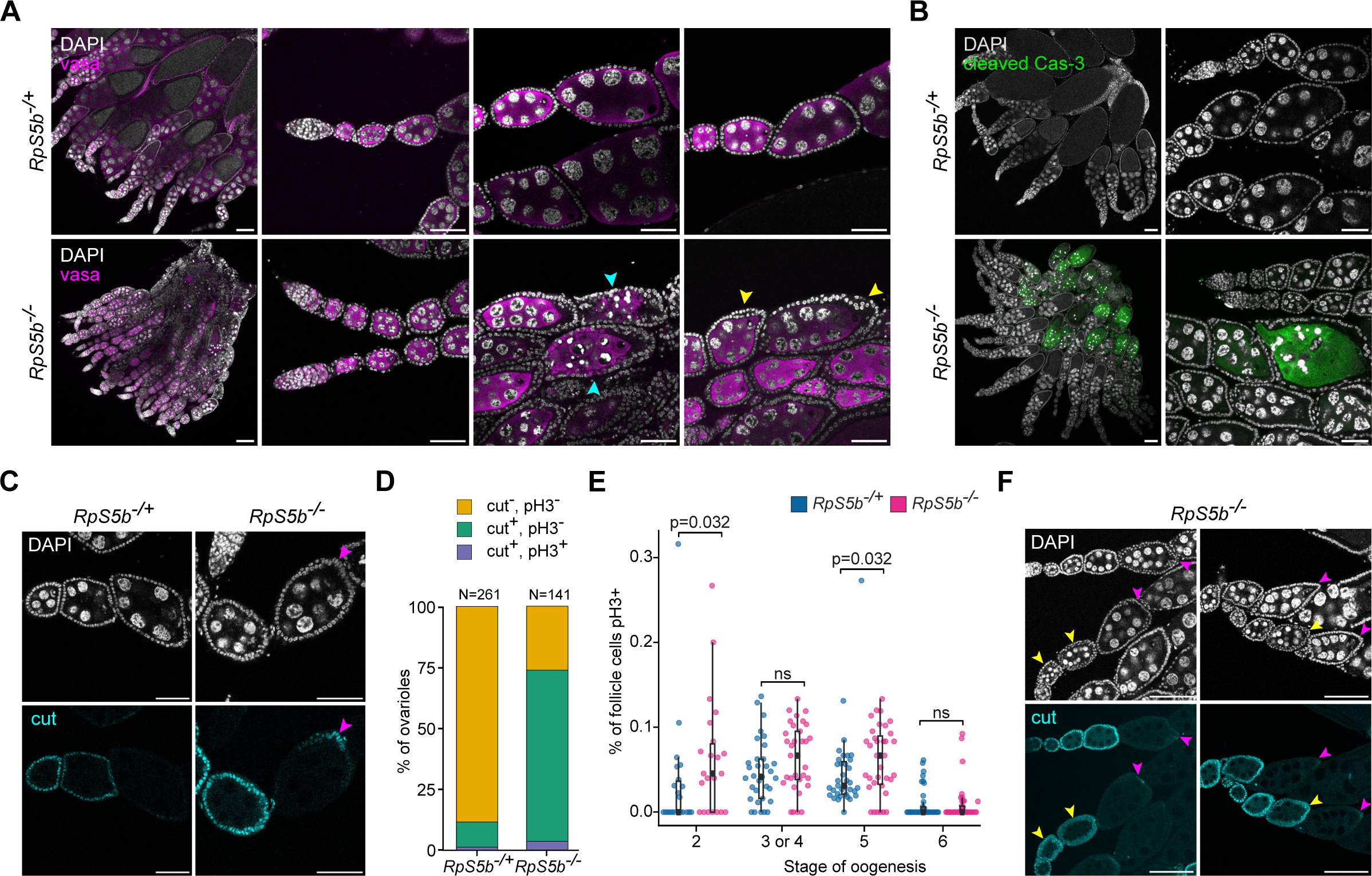
Characterisation of developmental defects in *RpS5b^-/-^* ovaries. **(A)** Representative confocal images of *RpS5b^-/+^* and *RpS5b^-/-^* ovaries, labelled with DAPI (DNA; grey) and vasa (germline; magenta). Cyan arrowheads indicate egg chambers with dying germ cells; yellow arrowheads indicate multilayering of follicle cells. Scale bars = 100µm for left panels, 50µm for all other panels. (**B**) Representative confocal images of *RpS5b^-/+^* and *RpS5b^-/-^* ovaries, labelled with DAPI (grey) and Cleaved Caspase-3 (green). Scale bars = 100µm for left images, 50µm for right images. (**C**) Representative confocal images of *RpS5b^-/+^* and *RpS5b^-/-^* ovaries, labelled with DAPI (grey) and Cut (cyan). Pink arrowheads indicate follicle cells with Cut signal at stage 6. Scale bars = 50µm. (**D**) Quantification of ovarioles in *RpS5b^-/+^* and *RpS5b^-/-^* ovaries containing egg chambers with follicle cells positive for pH3 and/or Cut beyond stage 6. (**E**) Quantification of the percentage of follicle cells positive for pH3 in egg chambers at different stages of oogenesis in *RpS5b^-/+^* and *RpS5b^-/-^* ovaries. For stage 2, N=22 for each genotype; for stage 3 and 4, N=33 for each genotype; for stage 5, N=35 for each genotype; for stage 6, N=37 for each genotype. *P-*values from Mann Whitney U test (Wilcoxon rank sum test). **F.** *RpS5b^-/-^* ovaries labelled with DAPI (grey) and Cut (cyan). Pink arrowheads indicate follicle cells with Cut signal at stage 6. Yellow arrowheads indicate egg chambers with multilayered follicle cells before stage 6. Scale bars = 50µm.

### Notch signaling is patchily dysregulated in *RpS5b^-/-^* egg chambers, but does not account for follicle cell multilayering

Bidirectional signaling between the germline and soma is essential for coordinating the growth of the two tissues ^47–49^. For instance, during egg chamber development, germline-to-soma Notch signaling triggers follicle cells to switch from mitotic cycles to endoreplication (Figure S5C ^50–52^), with disruption or delay of Notch activation leading to extra divisions of follicle cells. To test whether Notch signaling is disrupted in *RpS5b^-/-^* ovaries, we took advantage of antibodies against Cut and Hindsight (Hnt), two downstream readouts of the activity of the Notch signaling pathway. Similar to what was observed in heterozygous controls, Hnt expression was activated in stage 6 egg chambers in *RpS5b^-/-^* ovaries. Consistent with this observation, the overall expression of Cut consistently ceased just prior to stage 6 in both heterozygous and homozygous mutant ovaries. However, in contrast to controls, Hnt expression was not present in all follicle cells in *RpS5b^-/-^* ovaries. Indeed, we observed that Hnt was frequently absent from small patches of follicle cells located either at the posterior or in the middle of egg chambers (Figure S5D). Similarly, Cut expression was often observed in patches of posteriorly located follicle cells beyond stage 6 (73.8% of *RpS5b^-/-^* ovarioles, 11.5% of *RpS5b^-/+^* control ovarioles; Figure 3C-D). Together with previous observations (Jang *et al*., 2021), these results suggest that Notch signaling and downstream transduction is activated at stage 6 in *RpS5b^-/-^* ovaries, although dysregulation is frequently observed in small patches of cells.

As Notch signaling triggers the mitotic-to-endocycle switch, we tested whether this patchy dysregulation in *RpS5b^-/-^* egg chambers may cause an increase in follicle cell divisions. Antibody staining against the mitotic marker phospho-histone-3 (pH3) revealed that the number of ovarioles containing a stage 6 egg chamber with any pH3-positive follicle cells did not significantly differ between heterozygotes and homozygotes (Figure 3E). Additionally, DAPI staining and pre-rRNA single molecule Fluorescence In Situ Hybrydisation (smFISH) analyses demonstrated that the increase in nuclei and nucleoli volume from stage 5 to stage 7, proxies for the mitotic-to-endocycle switch, were observed in both *RpS5b^-/+^* and *RpS5b^-/-^* (Figure S5E). Follicle cells expressing Cut beyond stage 6 did not appear to have smaller nuclei than their Cut-negative neighbours. Together, these results suggest that the switch from mitosis to endocycling is successfully triggered in *RpS5b^-/-^* follicle cells, and that the multilayering phenotype is not a consequence of additional mitotic divisions post-stage 6 egg chambers. Interestingly, a small increase in the number of mitotic divisions, together with multilayering of follicle cells, was observed in *RpS5b^-/-^* egg chambers prior to the Notch-dependent switch at stage 6 (Figure 3E-F).

### *RpS5b^-/-^* germ cell death is independent of *p53* and *Xrp1* and resembles the mid-oogenesis checkpoint

Next we aimed to genetically dissect the pathways involved in the germ cell death induced by the lack of RpS5b. In mammals, impaired ribosome biogenesis leads to accumulation of the 5S RNP, which can bind to the E3 ubiquitin ligase MDM2 and prevent the ubiquitination and degradation of p53, thereby triggering apoptosis ^53,54^. Additionally, *p53* mutations have been shown to ameliorate the phenotypes of mouse models of ribosomopathies ^55,56^, therefore we recombined *RpS5b* mutant alleles with mutations in the *p53* gene ^57,58^. Our analyses revealed that germ cell death was not reduced in *p53^-/-^ RpS5b^-/-^* double mutants when compared to the *RpS5b^-/-^* single mutant (Figure 4A). Indeed, no eggs were produced in *p53^-/-^ RpS5b^-/-^-* double mutants, suggesting that p53 stabilisation is not the cause of the germ cell death induced by the lack of RpS5b.

**Figure 4.**
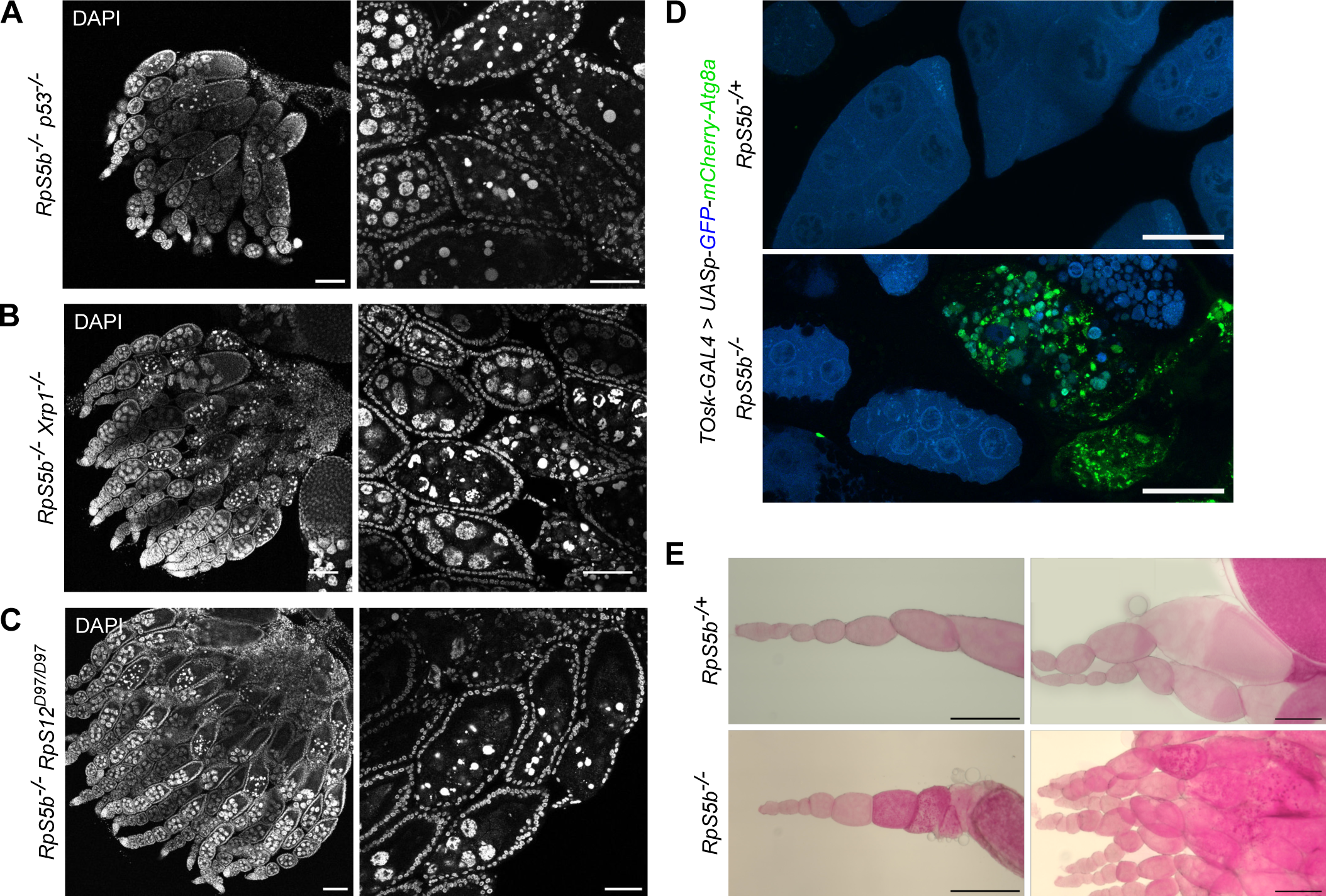
Germ cell death in *RpS5b^-/-^* ovaries is triggered independently of *p53* and *Xrp1*, and is associated with increased autophagy and premature glycogen accumulation. (**A-C**) Representative confocal images of (A) *RpS5b^-/-^ p53^-/-^*, (B) *RpS5b^-/-^ Xrp1^-/-^*, (C) and *RpS5b^-/-^ RpS12^D97/D97^* ovaries, labelled with DAPI (DNA). Scale bars = 100µm for left images, 50µm for right images. (**D**) Representative confocal images of *RpS5b^-/+^* and *RpS5b^-/-^* ovaries expressing the autophagy reporter *UASp-GFP-mCherry-Atg8a* (DeVorkin and Gorski, 2014) in germ cells, using the *TOsk-GAL4* driver (ElMaghraby *et al.*, 2022). GFP fluorescence is quenched by the acidic hydrolases in autolysosomes, which are marked by mCherry signal (DeVorkin and Gorski, 2014). GFP in blue, mCherry in green. Scale bars = 50µm. (**E**) Representative brightfield microscopy images of *RpS5b^-/+^* and *RpS5b^-/-^* ovaries stained with Periodic Acid Schiff’s reagent. Pink staining shows glycogen accumulation. Scale bars = 100µm.

Mutations in *Xrp1*, a gene encoding a bZip transcription factor, have been shown to suppress the apoptosis observed in *minute* tissues, by preventing activation of the integrated stress response (ISR) ^59–65^. To test whether *Xrp1* is required for promoting germ cell death in *RpS5b^-/-^*, we recombined a well-characterised *Xrp1* allele, *Xrp1^M2-73^* ^62,65^, with two *RpS5b* alleles. Our results revealed that *Xrp1^-/-^ RpS5b^-/-^* double mutant ovaries are phenotypically indistinguishable from single *RpS5b^-/-^* mutants (Figure 4B). Additionally, a specific point mutation in *RpS12* that prevents the activation of *Xrp1* in *minute* backgrounds (*RpS12^D97/D97^* ^61^) had no effect on the *RpS5b^-/-^* phenotype (Figure 4C). Altogether, these results indicate that Xrp1-dependent activation of the ISR is not involved in the germ cell death observed in *RpS5b^-/-^* ovaries.

Given that *RpS5b^-/-^*-induced death does not rely on either p53 or Xrp1, and based on the stereotyped development stage at which egg chambers collapse, we sought to test whether the *RpS5b^-/-^* phenotype may be related to the mid-oogenesis checkpoint. This well-characterised developmental checkpoint occurs before developing egg chambers enter into the energetically-expensive vitellogenesis phase, and has been most-studied in response to starvation ^66–68^. The distinguishing features of the mid-oogenesis checkpoint are reliance on activity of both the caspase Dcp-1 and autophagy, whilst being independent of p53 stabilisation ^69–71^. Additionally, the mid-oogenesis checkpoint occurs specifically at stage 8 of oogenesis, coinciding with a dip in the expression of apoptotic inhibitor DIAP-1 ^72^. As the *RpS5b^-/-^*-induced death occurs at stage 8 of oogenesis, is not rescued by p53 mutation, and involves cleavage of effector caspases, we decided to investigate autophagy in *RpS5b^-/-^* ovaries. Expression of the *UASp-GFP-mCherry-Atg8a* reporter ^73^ revealed an accumulation of autolysosomes in dying germ cells, demonstrating the occurrence of autophagy (Figure 4D). Together with the other characteristic features, this suggests that death of *RpS5b^-/-^* germ cells could be due to the activation of the mid-oogenesis checkpoint.

### Loss of RpS5b leads to ectopic TORC1 activation and premature metabolic remodelling

Soon after the mid-oogenesis checkpoint, germ cell metabolism is remodelled to a state of respiratory quiescence, which allows the accumulation of glycogen, an essential nutrient for embryogenesis ^74^. To examine whether germline metabolism was dysregulated in *RpS5b^-/-^*, we used Periodic Acid Schiff’s reagent assay to visualise glycogen accumulation during oogenesis. In contrast to *RpS5b^-/+^* heterozygotes and wild-type ovaries, in which an increase in glycogen accumulation is only observed from stage 12 onwards, the onset of glycogen accumulation in *RpS5b^-/-^* mutants occurred from stage 5-6 of oogenesis (Figure 4E).

Interestingly, this early glycogen accumulation was also observed in germline knockdowns of components of the insulin signaling pathway, as well as in response to starvation ^74^. The evolutionarily conserved Tor kinase integrates a variety of nutritional inputs, including insulin signaling and amino acids, to coordinate growth and metabolism (Figure 5A ^75^). To investigate whether Tor signaling is disrupted in *RpS5b^-/-^* ovaries, we took advantage of phospho-specific antibodies against p-RpS6 (pS6) and p-4E-BP, two downstream readouts of TORC1 activity ^76^. In wild type and heterozygous mutants, p-4E-BP is detected throughout germline development, increasing from the end of the germarium onwards (Figure 5B; ^77^), while pS6 is exclusively observed in germ cells, in a short developmental window within the germarium corresponding to the 2-8 cell cysts stages ^20^. In *RpS5b^-/-^* mutants, we did not observe any changes in p-4E-BP staining. In striking contrast, strong pS6 signal was ectopically observed throughout egg chamber development in *RpS5b^-/-^* mutants, from region 2b until egg chambers collapsed (Figure 5C). Furthermore, mutants also displayed strong pS6 signal in somatic follicle cells, frequently displaying a “salt-and-pepper” pattern. Notably, follicle stem cells, located at the end of the germarium, were pS6+ in 42.7% of *RpS5b^-/-^* ovarioles, in contrast to 0.3% in *RpS5b^-/+^* controls (Figure 5D-E). This ectopic activation of TORC1 in 16-cell-cysts and follicle stem cells is the earliest abnormality we observed, occurring many developmental stages before the somatic multilayering and the egg chamber death phenotypes.

**Figure 5.**
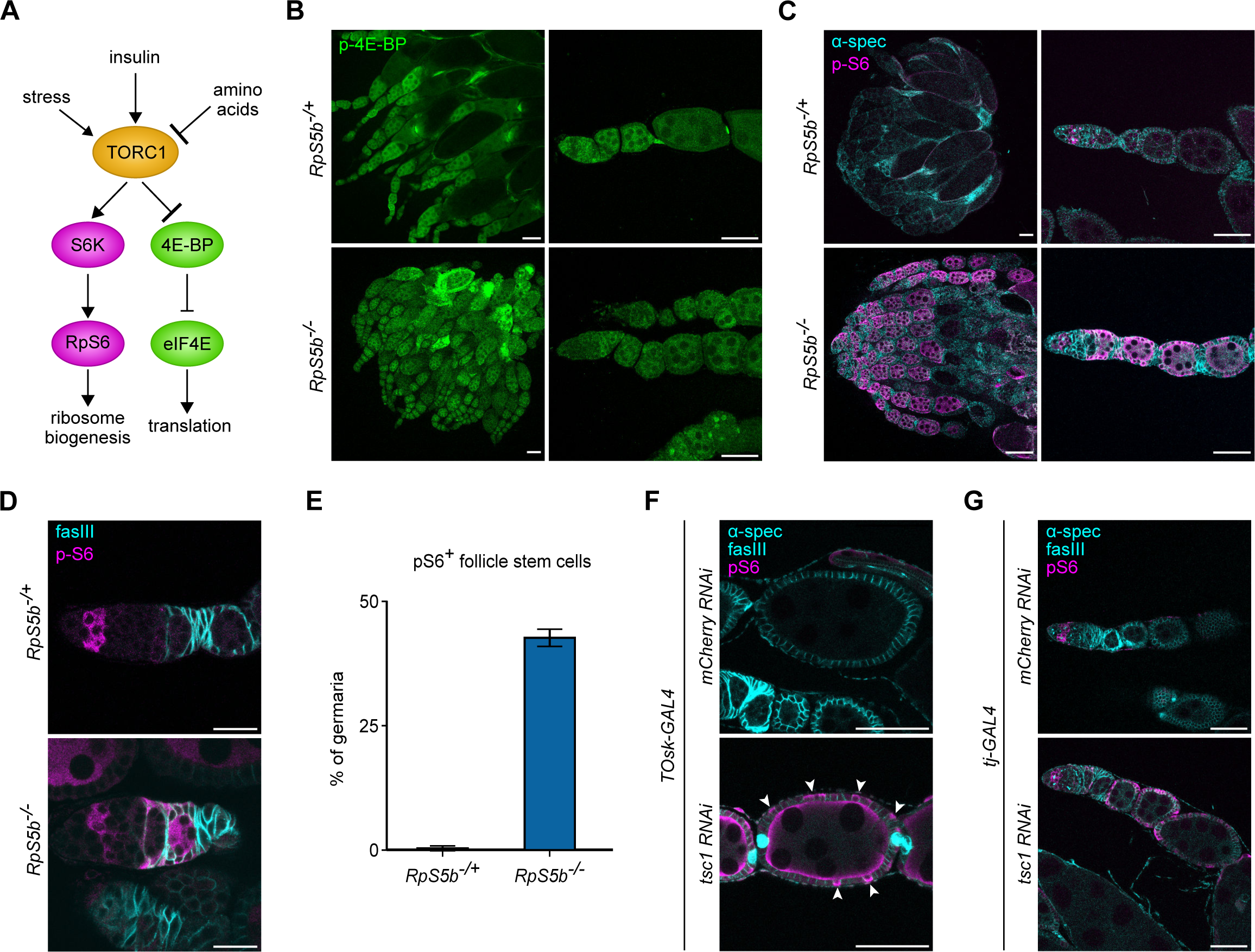
Loss of RpS5b triggers TORC1 activation in germline and soma. (A) Schematic showing simplified TORC1 regulation of ribosome biogenesis and translation. (**B-C**) Representative confocal images of *RpS5b^-/+^* and *RpS5b^-/-^* ovaries, labelled with (B) p-4E-BP (green), or (C) α-spectrin (spectrosomes and fusomes, cyan) and p-RpS6 (pS6, magenta). Scale bars = 100µm for left images, 50µm for right images. (**D**) Representative confocal images of *RpS5b^-/+^* and *RpS5b^-/-^* germaria, labelled with fasciclin-III (follicle cell membranes, cyan) and pS6 (magenta). Scale bars = 20µm. (**E**) Quantification of *RpS5b^-/+^* (N=217) and *RpS5b^-/-^* (N=344) germaria containing pS6-positive follicle stem cells. Error bars show standard deviation across three replicates. (**F-G**) Representative confocal images of (F) *TOsk-GAL4 > mCherry RNAi* and *TOsk-GAL4 > tsc1 RNAi* ovaries, and (G) *Tj-GAL4 > mCherry RNAi* and *Tj-GAL4 > tsc1 RNAi* ovaries, labelled with DAPI (DNA, grey), α-spectrin and fasciclin-III (spectrosomes, fusomes and follicle cell membranes, cyan), and pS6 (magenta). In (F), white arrowheads indicate pS6-positive follicle cells. Scale bars = 50μm.

To test whether TORC1 activity in somatic cells is regulated in a cell-autonomous manner or is a consequence of the ectopic activation of TORC1 in germ cells, we used the germline-specific *TOsk-GAL4* driver to knockdown *tsc1*, a well-described repressor of TORC1 activity ^78,79^. As expected, germline knockdown of *tsc1* led to the ectopic activation of TORC1 in germ cells (Figure 5F). However, remarkably, we also observed ectopic TORC1 activation in the neighbouring somatic cells, in a “salt-and-pepper” pattern of pS6 signal, reminiscent of that observed in *RpS5b^-/-^* egg chambers. Given that RpS5b is exclusively required in germ cells, these results suggest that the *RpS5b^-/-^*-induced germline TORC1 activation triggers non-autonomous Tor activation in the soma, revealing a novel layer of inter-tissue communication. Interestingly, this mechanism seems to be unidirectional, as ectopic activation of TORC1 in somatic cells using the *Tj-Gal4* driver to knockdown *tsc1* did not induce a similar response in germ cells (Figure 5G).

### Disrupting canonical ribosomal proteins or ribosome biogenesis during egg chamber development recapitulates the *RpS5b^-/-^* phenotype

Given that *RpS5b* is predominantly expressed in the germline from stage 2b onwards, we examined whether the depletion of canonical ribosome proteins at the same developmental stage could induce similar phenotypes to those observed in *RpS5b**^-/-^*** mutants. To do this, we used the *TOsk-GAL4* driver, which is specifically activated from stage 2 onwards ^78^, to knockdown other RPs. Similar to what was observed in *RpS5b* mutants, individual germline knockdown of *RpS17*, *RpL35* and *RpS18* caused ectopic S6 phosphorylation and egg chamber death (Figure 6A). Moreover, germline-specific RNAi was sufficient to induce somatic S6 phosphorylation, follicle cell multilayering, and Notch dysregulation (Figure 6B). Additionally, we also observed premature glycogen accumulation (Figure S6A) as in *RpS5b**^-/-^***.

**Figure 6.**
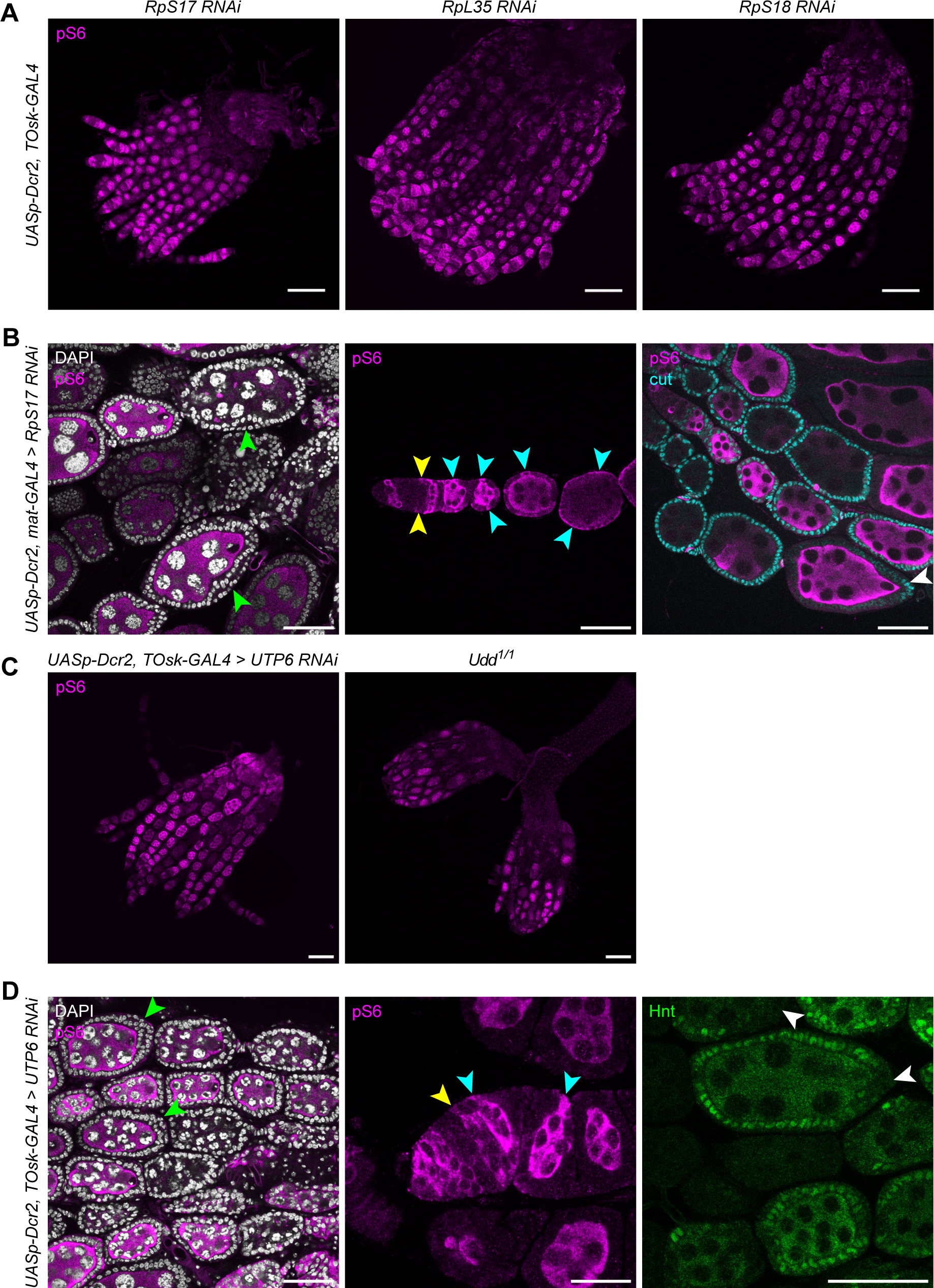
Disruption of canonical ribosomal proteins or ribosome biogenesis factors during egg chamber development mimics the *RpS5b^-/-^* phenotype. (**A**) Representative confocal images of *TOsk-GAL4 > RpS17 RNAi*, *RpL35 RNAi* and *RpS18 RNAi* ovaries, labelled with pS6 (magenta). Scale bars = 100µm. (**B**) Representative confocal images of *Mat-tub-GAL4, UASp-Dcr2 > RpS17 RNAi* ovaries, labelled with DAPI (DNA, grey), pS6 (magenta) and Cut (cyan). Green arrowheads indicate multilayering of follicle cells; cyan arrowheads indicate pS6-positive follicle cells; yellow arrowheads indicate pS6-positive follicle stem cells; white arrowheads indicate Cut-positive follicle cells at stage 6. Scale bars = 50µm. (**C**) Representative confocal images of *TOsk-GAL4 > UTP6 RNAi* and *Udd^1/1^* ovaries, labelled with pS6 (magenta). Scale bars = 50µm. (**D**) Representative confocal images of *TOsk-GAL4 > UTP6 RNAi* ovaries, labelled with DAPI (DNA, grey), pS6 (magenta) and Hnt (green). Green arrowheads indicate multilayering of follicle cells; cyan arrowheads indicate pS6-positive follicle cells; yellow arrowheads indicate pS6-positive follicle stem cells; white arrowheads indicate Hnt-negative follicle cells at stage 6. Scale bars = 50µm for left and right images; 20µm for middle image.

To deplete ribosome levels through an alternative mechanism, we perturbed ribosome biogenesis (RiBi) by using either mutants or germline-specific knockdowns. Udd promotes RNA polymerase I-mediated pre-rRNA transcription ^19^, while UTP6 is involved in the cleavage of the 47S pre-rRNA ^80^. Ovaries of both *udd^1/1^* hypomorphic mutants and germline-specific *UTP6* knockdowns presented all the hallmark phenotypes observed in *RpS5b* mutants: germline and somatic S6 phosphorylation, germ cell death, follicle cell multilayering, and Notch dysregulation (Figure 6C-D). Furthermore, ovaries of *pelota^PB60/PA13^* hypomorphic mutants, a key ribosome recycling factor, display a similar phenotype including TORC1 activation and collapse of the majority of egg chambers by mid-oogenesis (Figure S6B). Altogether, these results suggest that the *RpS5b^-/-^* phenotype is similar to that triggered by insufficient functional ribosomes in germ cells during egg chamber development.

### RpS5a and RpS5b are functionally interchangeable

It has been previously proposed that ribosomes containing RpS5a and RpS5b may be functionally different, with RpS5b-containing ribosomes providing “specialised” functions and translating distinct subsets of mRNAs ^43^. To definitively test whether RpS5a and RpS5b proteins are functionally distinct, we used CRISPR/Cas9 and homology-directed repair (HDR) to seamlessly replace the entire coding sequence of *RpS5b* with the coding sequence of *RpS5a*, at the endogenous locus. This was engineered so that the *RpS5b* 5’- and 3’-UTRs, promoter, and all surrounding regulatory regions were intact, ensuring that the *RpS5a* coding region would be expressed and regulated in the same way as RpS5b (*RpS5b^RpS5a/RpS5a^*; Figure 7A). The reciprocal edit was also independently carried out to generate flies in which the coding sequence of *RpS5a* was replaced by that of *RpS5b*, whilst maintaining all *RpS5a* regulatory sequences. Of note, the introns of *RpS5a* encode 14 H/ACA box snoRNAs ^81^, and therefore, we engineered a strategy to maintain the *RpS5a* intron sequences while replacing most of the coding region (*RpS5a^RpS5b/RpS5b^*; Figure 7B and S7A; see Materials and Methods).

**Figure 7.**
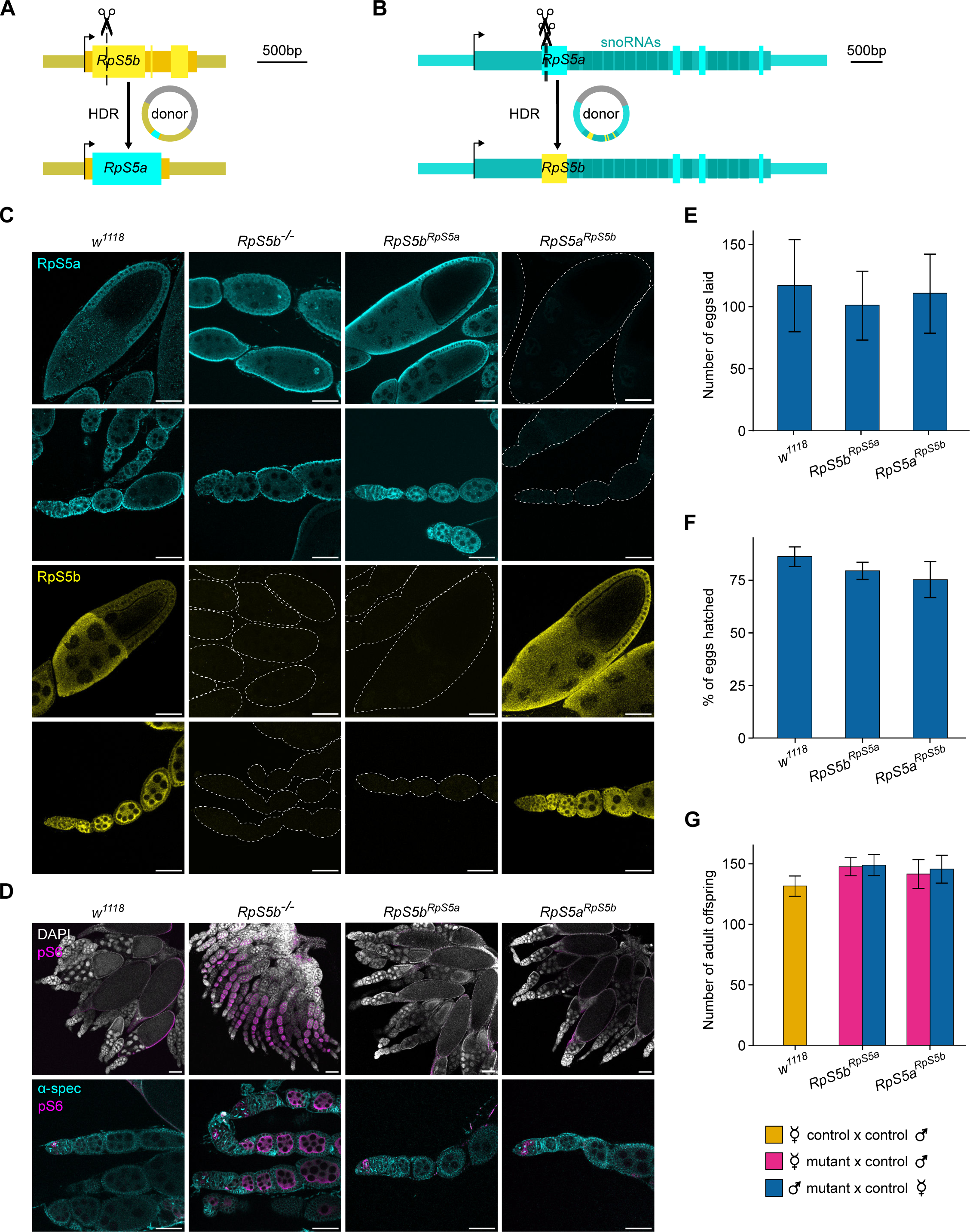
RpS5 paralogues are functionally interchangeable. (**A-B**) Schematic illustrating CRISPR-Cas9-based strategy used to replace coding sequence of (A) *RpS5b* with that of *RpS5a*, and (B) *RpS5a* with that of *RpS5b*. Sequences encoding snoRNAs in the RpS5a introns are labelled in darker blue. (**C**) Representative confocal images of *w^1118^* (controls), *RpS5b^-/-^*, *RpS5b^RpS5a/RpS5a^*, and *RpS5a^RpS5b/RpS5b^* ovaries, labelled with RpS5a (cyan) and RpS5b (yellow). White dotted lines show the outline of egg chambers when little/no signal was detected. Scale bars = 50µm. (**D**) Representative confocal images of *w^1118^* (controls), *RpS5b^-/-^*, *RpS5b^RpS5a/RpS5a^*, and *RpS5a^RpS5b/RpS5b^* ovaries, labelled with DAPI (grey), α-spectrin (spectrosomes and fusomes, cyan), and pS6 (magenta). Scale bars = 100µm for top row images, 50µm for bottom row images. (**E**) Quantification of egg laying for *w^1118^*, *RpS5b^RpS5a/RpS5a^*, and *RpS5a^RpS5b/RpS5b^* females (results for three biological replicates per genotype); error bars show standard deviation. (**F**) Quantification of egg hatching (same eggs as **E**; results for three biological replicates per genotype); error bars show standard deviation. (**G**) Quantification of adult offspring (results for three biological replicates per sex and genotype, for independent crosses from those in **E** and **F**); error bars show standard deviation.

First, we validated the stocks by using immunofluorescence to characterise the expression of RpS5a and RpS5b (Figure 7C). In *RpS5b^RpS5a/RpS5a^* ovaries, RpS5a was highly expressed in both the germline and soma whilst RpS5b was absent, confirming that the *RpS5a* CDS was indeed being expressed in the pattern of wild-type RpS5b, in addition to the endogenous RpS5a expression. Conversely, RpS5a was not detected in *RpS5a^RpS5b/RpS5b^* ovaries, whilst RpS5b was highly expressed in both the germline and the soma, confirming that RpS5b is expressed in the pattern of wild-type RpS5a, in addition to the endogenous RpS5b expression. Importantly, both edited stocks were viable as homozygotes, with no obvious phenotypic changes being observed. Moreover, the comparison of *RpS5a^RpS5b/RpS5b^* flies with wild type *w^1118^* flies and the classic *minute RpS5a^-/+^* revealed that *RpS5a^RpS5b/RpS5b^* flies displayed wild type bristles, demonstrating that RpS5b can seamlessly replace RpS5a with regards to the *minute* phenotype (Figure S7B).

We next characterised these flies in terms of ovary morphology and germline development. Upon dissection, both *RpS5b^RpS5a/RpS5a^* and *RpS5a^RpS5b/RpS5b^* adult ovaries appeared wild type, with egg chambers progressing through all stages of development to produce mature eggs. Immunostaining revealed that both *RpS5b^RpS5a/RpS5a^* and *RpS5a^RpS5b/RpS5b^* ovaries were also wild type with respect to S6 phosphorylation, with pS6 only being observed in the 2-8 cell cysts and being absent from any later stages of oogenesis (Figure 7D). Indeed, we did not observe any difference in either the number of eggs laid or the proportion of those eggs hatching between *RpS5b^RpS5a/RpS5a^*, *RpS5a^RpS5b/RpS5b^* and control *w^1118^* females (Figure 7E-F). Moreover, independent reciprocal crosses showed that there was also no difference in the overall number of adult offspring produced by *RpS5b^RpS5a/RpS5a^*, *RpS5a^RpS5b/RpS5b^* and *w^1118^* females or males, when crossed to control *w^1118^* flies (Figure 7G).

The lack of detectable phenotype in *RpS5b^RpS5a/RpS5a^* flies and *RpS5a^RpS5b/RpS5b^* flies shows that, at least under laboratory conditions, RpS5a and RpS5b proteins are functionally equivalent. We found no evidence to suggest RpS5b has a different function to RpS5a. Therefore, we conclude that the phenotypic abnormalities observed in *RpS5b^-/-^* mutants are the result of insufficient functional ribosomes being assembled in the germline from mid-oogenesis onwards, as RpS5b is the predominantly expressed RpS5 paralogue in that developmental window.

## Discussion

Recent work uncovering the heterogeneity in ribosome composition in a diverse array of species and tissues has led to the hypothesis that such so called “specialised ribosomes” may elicit qualitative changes in translation capacities in particular cell types or development stages. Our evolutionary analysis revealed that RP genes are frequently duplicated, potentially fuelling the arisal of new diversity. However, our results indicate that only a very small subset of duplicates persist for long time periods or show any evidence of transcription. Focusing on those that are retained through evolution and transcriptionally competent, we performed a systematic functional analysis in *D. melanogaster* by generating loss-of-function mutants for 11 RP paralogues, seven of which are expressed almost exclusively in the testes. 10 of these paralogues were not required for development or fertility, and mutants displayed no obvious phenotype under lab conditions. The only paralogue for which mutations led to phenotypic abnormalities, *RpS5b*, was shown to be functionally redundant with its canonical parental protein RpS5a. Indeed, we demonstrate that phenotypic abnormalities were a consequence of differences in their expression patterns and resulted from a deficit in the total levels of RpS5 - rather than any protein or function specialisation. Therefore, the systematic study of the RP paralogues in *D. melanogaster*, the first of the kind, provides little-to-no evidence to support the existence of “specialised ribosomes”. Instead, this work emphasises the importance of matching ribosome levels to the tissue requirements, and in turn uncovered a previously unknown inter-tissue signalling response to ribosome insufficiency.

Our results reveal that the majority of duplicates identified in the *Drosophila* genus arose by retroposition, as has been the case in mammals ^82–84^. In that clade, the high frequency of retroposition of RP genes is thought to be due to their high expression levels, and previous analysis has found that almost 20% of pseudogenes in the human, mouse and macaque genomes originated from RP genes ^83^. Similar to what we observed in *Drosophila*, the majority of duplicates identified in mammalian genomes were evolutionarily recent, suggesting that RP duplicates are lost at a quick pace. As the majority of retrogenes are not expressed ^85,86^, selection against dominant-negative effects elicited by duplicated RP genes - for instance, the perturbation of the stoichiometries of ribosomal protein required for proper ribosome biogenesis ^87^ - seems unlikely to account for the turnover. Instead, untranscribed duplicates are most likely to evolve neutrally, gradually accumulating mutations, transposable element insertions, or being lost via recombination. Indeed, comprehensive analysis of human and *Drosophila* pseudogenes revealed an enrichment of pseudogenes in areas of low recombination ^83^.

One open question is why a number of RP paralogues have been maintained for millions of years during *Drosophila* evolution ^27^, despite seemingly not being required for viability or fertility. In the case of the *RpS5* paralogues, the retention of both paralogues most likely represents an example of Duplication-Degeneration-Complementation (DDC; ^88,89^, whereby functionally redundant paralogues can become essential and conserved over long periods if complementary loss of expression domains occurs, via mutation of *cis*-regulatory elements. A plausible scenario is that, prior to duplication, the ancestral *RpS5a* was highly expressed in all tissues. Immediately after the DNA-mediated duplication event that gave rise to *RpS5b*, there would likely have been redundancy between the two genes. A later change in the cis-regulatory elements led to a reduction of *RpS5a* expression in the germline. At this point, germline *RpS5b* expression became essential, therefore being maintained by selective pressure.

The retention of the remaining RP paralogues is more puzzling, although the majority are specifically expressed in the *D. melanogaster* testis. Across a variety of species, the testis is the tissue with the largest number of genes expressed ^90,91^, with either a permissive chromatin environment or the linkage of DNA repair to transcriptional scanning hypothesised to underlie this widespread transcription ^91–93^. If RP retrocopies were expressed in the testis, for instance due to insertion close to an existing promoter, they could then be influenced by selection and potentially maintained. Indeed, human studies have identified a number of retrocopies that are expressed in the testis and show signatures of purifying selection ^86^. Perhaps, despite these RP paralogues not being required in laboratory conditions, they may provide a slight reproductive advantage by ensuring the robustness of spermatogenesis under specific environmental conditions.

The majority of the RP paralogues we investigated here have been previously shown to assemble into translating ribosomes in germ cells, albeit often comprising small populations of the total ribosome pool ^24^. Our results suggest that, at least under laboratory conditions, the ribosomal heterogeneity provided by such RP paralogues is not required for normal somatic and germline development, viability or fertility. Instead, our results indicate that producing sufficient ribosomes is a key requirement for oogenesis. This is not surprising, as nurse cells must produce all proteins and ribosomes to be deposited into the oocyte and sustain embryogenesis until zygotic genome activation ^94^. However, in addition to restricting global levels of translation, reduced ribosome levels may also particularly decrease the translation of specific transcripts ^95^. Data gathered in yeast revealed that the most highly translated transcripts were the most dramatically affected by a decrease in ribosome levels ^96^. This suggests that *RpS5b^-/-^* egg chambers may also be deficient in specific proteins, such as the yolk proteins (as noted by Jang *et al* ^44^, microtubule components, and other translational machinery which are all amongst the most highly expressed factors in mature oocytes and are essential for embryogenesis ^97^.

Thanks to the analysis of *RpS5b* mutants, we were able to uncover the existence of a mechanism coordinating TORC1 activation between the germline and the surrounding somatic follicle cells. While it is well established that inter-tissue communication is required to coordinate growth between the germline and the soma ^47,48,52,98,99^, TORC1 coordination does not seem to be mediated by previously known signalling pathways. Intriguingly, our results reveal that this coordination is unidirectional, from the germline to the soma. In *C. elegans* neurons, TORC1 activity has been shown to promote the release of insulin-peptides ^100^, promoting the activation of TORC1 in distant tissues. However, it is unlikely that the coordination observed in the *Drosophila* ovary relies on the release of systemic signals such as insulin-like peptides, as our results suggest that the non-autonomous activation in ovarian somatic follicle cells relies on direct contact with germ cells. One possibility is that the transducing signal could be directly shared between these cells via gap junctions ^101^. Innexins, the structural component of gap-junctions, are expressed in nurse cells and oocytes, and have been shown to be required for the early stages of germ cell differentiation ^102–104^. In mice, it has been shown that amino acids are transferred from somatic cells to growing oocytes via gap junctions ^105^. While it remains to be determined how TORC1 activation in the germline can be transduced into somatic cells, it is tempting to speculate that such a mechanism is important for synchronising the growth of both tissues during egg chamber development.

In the *Drosophila* somatic tissues studied thus far, a deficit in ribosome production triggers the activation of the ISR, via Xrp1, RpS12 and PERK ^61,62,65,106–108^. ISR activation is followed by global suppression of translation initiation, reduced ribosome biogenesis, and concomitant upregulation of genes involved in autophagy and protein folding ^109^. This is thought to allow proteostasis to be restored and to promote cell survival, with apoptosis being triggered only by persistent ISR activation ^107^. Intriguingly, this is diametrically opposite to the response observed when developing germ cells are depleted of ribosomes: we revealed that a deficit in ribosomes in the germline leads to acute TORC1 activation, which is associated with an increase in both ribosome biogenesis and global translation ^44^. Such contrasting responses raise key questions about how tissues experiencing ribosome insufficiency determine which pathway to employ. One possibility is that the outcome may depend upon the severity of the ribosome insufficiency. For instance, the ISR has mainly been studied in *minute* tissues heterozygous for RP mutants, which present a 0-30% reduction in ribosomes compared to wild-type ^64^. Nevertheless, knockdown of RPs in wing disc cells also leads to activation of ISR ^62^. Alternatively, cellular response may depend on translational requirements, which are likely to vary from tissue to tissue, as well as during development and homeostasis. In this context, the female germline is particularly known for its growth requirements, due to the demands of sustaining the early stages of embryogenesis.

Importantly, the balance between ISR activation and growth is also likely to play a key role in human ribosomopathies. For instance, persistent activation of the ISR restricts growth and triggers cell death via *p53* ^110^, which can result in developmental phenotypes such as skeletal deformities and severe anaemia. However, this response places a strong selective pressure on affected tissues to produce sufficient ribosomes. Indeed, most ribosomopathies are associated with cancer, likely due to these conditions favouring the proliferation of cells with compensatory mutations that bypass ribosome quality control, or cellular surveillance mechanisms such as p53 ^111–113^. Manipulation of these opposing forces could be exploited to treat ribosomopathies. For example, overactivation of TORC1 in brain organoids modelling patient mutations in the ribosome biogenesis factor *AIRIM* rescued the translation defect and associated neurodevelopmental phenotype ^114^; whilst early trials using L-leucine to activate TORC1 have shown some promise in improving growth and haematopoiesis in Diamond-Blackfan anaemia patients ^115,116^. In this context, our findings highlight that understanding the unique ribosome biogenesis and translation demands of different tissues and developmental stages, in addition to how these are sensed, will be essential to make use of potential new therapies without driving cancer.

## Resource availability

### Lead contact

Further information and requests for resources or reagents should be directed to the lead contact, Felipe Karam Teixeira.

### Materials availability

All transgenic *Drosophila* lines generated in this study are available upon request.

### Data and code availability

Custom code used in this study is deposited at Github (https://github.com/d-gebert/RP_blast). Any additional information required to reanalyse the data reported in this study is available upon request.

## Supporting information

Key resource table

## Acknowledgements

The authors thank Aurelio Teleman, Paul Lasko, Julie Aspden, Ruth Lehmann and Daniel St Johnston for antibodies; the Vienna Drosophila Resource Center, the Bloomington Drosophila Stock Center, and Eric Lai, Nicholas Baker, Paul Lasko, Michael Buszczak, Golnar Kolahgar, and Ruth Lehmann for fly reagents; and Tamsin Samuels for discussion and comments on the manuscript. The authors gratefully acknowledge the Department of Genetics Fly Facility, and Dr Ian Clark, Dr Antonina J. Kruppa, Dr Ben Sutcliffe, and Dr Jonathan D. Howe from the Department of Genetics imaging facility for their support and assistance in this work and thank the Wellcome Trust for a strategic award (105602/Z/14/Z). KZAG is an MRC PhD student and is also supported by the Cambridge Philosophical Society and EMBO Bridging Fund. DG is supported by a Walter Benjamin Postdoctoral Fellowship from Deutsche Forschungsgemeinschaft (GE3407/1-1). CS was supported by a Stephen Johnson Research Bursary. FKT is a Wellcome Trust and Royal Society Sir Henry Dale Fellow (206257/Z/17/Z), an EMBO YIP Investigator (5025), and is supported by the Human Frontier Science Program (CDA-00032/2018). For the purpose of Open Access, the author has applied a CC BY public copyright licence to any Author Accepted Manuscript (AAM) version arising from this submission.

## Author contributions

KZAG and FKT conceived the idea and designed the experiments. KZAG, CS, and FKT performed the experiments. KZAG and DG performed bioinformatic analyses. KZAG and FKT wrote the manuscript, with input from all authors.

## Supplemental information

Document S1: Figures S1-S7, Tables S3-S5, legends for Tables S1-S2

Table S1 (excel file)

Tables S2 (excel file)

## Material and Methods

### Drosophila methods

Unless stated otherwise, stocks and crosses were maintained at 25°C on standard propionic food, prepared by Cambridge University Fly Facility.

To assess fertility, 10 mutant virgin females and 10 *w^1118^* males were allowed to mate and lay for 2 days at 25°C. Parents were then removed and the number of adult offspring was counted after 12 days. Crosses were carried out in triplicate and reciprocally. To assess egg laying and hatching, four females and four males were allowed to mate for 24 hours at 25°C. Flies were then flipped to a new vial of propionic food and allowed to lay for 24 hours. Adults were then removed and the number of eggs laid were counted. Vials were returned to 25°C and the number of hatched eggs was counted 24 hours later.

To induce recombination for clonal analysis, vials containing 3rd instar larvae were incubated at 37°C for one hour, then incubated at 34°C for 2 hours. Vials were then returned to 25°C until eclosion. All experiments were conducted in at least 2 biological and 2 technical replicates.

For embryo collections, 20-40 flies were placed overnight in cages with apple-agar plates and wet yeast paste. The following morning, adult flies were flipped to new yeasted apple-agar plates twice, each one hour apart, and these plates were discarded. Then, plates were flipped every two hours, and left to develop at 25°C for the desired length of time. To dechorionate embryos, plates were rinsed with 50% bleach (2.6% sodium hypochlorite) for 2 minutes. Embryos were then collected in a fine mesh, rinsed well with water, then transferred to PBS before being used for immunofluorescence or Western blot analyses.

### Immunofluorescence

Adult ovaries and testes were dissected in ice-cold PBS buffer and fixed in PBST (PBS with 0.2% Triton X-100) containing 4% Formaldehyde (Thermo Fisher Scientific) for 30 min. Fixed tissues were rinsed three times with PBST before incubation in blocking buffer (PBST with 1% BSA) for one hour at room temperature. Samples were then incubated with primary antibody diluted in blocking buffer overnight at 4°C. Samples were washed five times with PBST, then incubated with secondary antibodies diluted in blocking buffer overnight at 4°C. Samples were washed five times with PBST and mounted in VectaShield mounting medium with DAPI (Vector Laboratories). Fluorescent images were acquired on a Leica SP8 confocal microscope using 10x, 20x and 40x objectives. Images were processed using Fiji. All experiments were conducted in at least 2 biological and 2 technical replicates. Dechorionated embryos were incubated for 30 minutes in fixative (500μl 37% formaldehyde, 1.5ml PBS and 8ml heptane) whilst shaking. Embryos were washed 3 times in PBST, then hand-devitellised in PBS. Primary and secondary antibody staining, mounting and imaging were carried out as described for ovaries and testes. All experiments were conducted in at least 2 biological and 2 technical replicates.

### smFISH

Ovaries were dissected and fixed as described for IF. After 3× washes in PBST, samples were transferred to Wash buffer (2× saline sodium citrate (SSC), 10% deionised formamide in nuclease-free water) for 10 minutes at room temperature. Pre-rRNA smFISH probes (Table S3) were diluted in Hybridisation buffer (2× SSC, 10% deionised formamide, 20 mM vanadyl ribonucleoside complex, 0.1 mg/ml BSA, competitor (1:50 dilution of 5 mg/ml *E. coli* tRNA and 5 mg/ml salmon sperm ssDNA) in nuclease-free water). Ovaries were incubated in Hybridisation buffer at 37 °C overnight. Ovaries were washed three times for 15 minutes in Wash buffer. In the penultimate wash step, DAPI was added at 1:200. After washing, samples were mounted in VectaShield mounting medium and imaged as described above. Experiments were conducted in at least 2 biological and 2 technical replicates.

### Periodic Acid Schiff’s staining

To visualise glycogen accumulation, ovaries were dissected in ice-cold M3 media, then fixed in 4% formaldehyde in M3 media for 30 minutes. Samples were then washed three times with distilled water, incubated in periodic acid solution for 15 minutes, then washed four times with distilled water. Samples were then incubated in Schiff’s reagent for four minutes, then washed three times with distilled water. Finally, samples were washed twice with PBS and then mounted in PBS. Slides were imaged by brightfield microscopy at 5x and 10x using a Zeiss Axiophot. All experiments were conducted in at least 2 biological and 2 technical replicates.

### Western blot

20-30 pairs of ovaries or testes, or 50-100 embryos, were collected for each sample and homogenised in RIPA buffer (150mM NaCl, 1.0% NP-40, 0.5% deoxycholate, 0.1% SDS, 50mM Tris-Cl). After quantification by Bradford assay, Laemmli buffer with β-mercaptoethanol was added, and samples were incubated at 95 °C for 10 minutes. Samples were run on a Novex Value 16% Tris-Glycine Mini Protein Gel then wet transferred to a PVDF membrane. Blots were blocked with 5% skimmed milk in TBST (TBS with 1% Tween) for 1 h at room temperature. Primary antibody incubation was at 4 °C overnight, then blots were washed and incubated with secondary antibodies in TBSTS (TBST with 0.01% SDS) for 30 minutes at room temperature. Blots were imaged with a LI-COR Odyssey imager. All experiments were conducted in at least 2 biological and 2 technical replicates.

### CRISPR/Cas9

Guide RNAs were designed using flyCRISPR Optimal Target Finder ^117,118^. As previously described ^117^, oligos (listed in Table S4) were then annealed to each other, before ligation into the pU6-BbsI-gRNA plasmid, linearised with BbsI.

For mutagenesis, pairs of guide plasmids targeting a single gene were co-injected into *nos-cas9* eggs by the Cambridge University Fly Facility (*y[1] M{w[+mC]=nanos-Cas9.P}ZH-2A w[*]* eggs for all genes except *RpS14b*, where *y[1] sc[1] v[1];; {y[+t7.7] v[+t1.8]=nanos-Cas9}attp2* was used). Screening was carried out by PCR and Sanger sequencing (Genewiz), and balanced stocks were established.

To produce the donor for replacing the *RpS5b* coding sequence with that of *RpS5a*, RNA was extracted from *w^1118^* flies and RT-PCR used to amplify the CDS of *RpS5a*. DNA was extracted from *w^1118^* flies and 1.7kb homology arms including the 5’ and 3’UTRs of *RpS5b* were amplified by PCR. The CDS and homology arms were Gibson assembled into the *pBluescript II SK(+)* plasmid, digested with HindIII. NEBuilder Assembly Tool was used to design primers with overhangs for Gibson Assembly. To produce the donor for replacing the *RpS5a* coding sequence with that of *RpS5b*, we designed a donor plasmid such that only the *RpS5a* exons were replaced with the *RpS5b* coding sequences, whilst the introns of *RpS5a* were unaltered, and 1.7kb homology arms were included. This plasmid was synthesised by Genewiz. In both cases, the CRISPR guides had been designed to target the divergent N-terminus of the coding sequences, so the PAMs were not present in the donor sequences.

For targeting *RpS5b*, the appropriate donor plasmid and guide plasmids targeting *RpS5b* and *ebony* were injected into *y[1] M{w[+mC]=nanos-Cas9.P}ZH-2A w[*]* eggs by the Cambridge University Fly Facility. A co-CRISPR strategy was used ^119^, with *ebony* F^1^ being used to establish stocks and screened by PCR. For targeting *RpS5a*, the appropriate donor plasmid and guide plasmids targeting *RpS5b* and *white* were injected into *y[1] sc[1] v[1];; {y[+t7.7] v[+t1.8]=nanos-Cas9}attp2* eggs by the Cambridge University Fly Facility. *White* F^1^ individuals were used to establish stocks and screened by PCR and Sanger sequencing.

Targeting of *RpS5b* resulted in the complete replacement of the *RpS5b* coding sequence with that of *RpS5a*. Targeting of *RpS5a* resulted in a chimaeric RpS5 protein, where all but 6 amino acids of RpS5a had been replaced by the sequence of RpS5b (Figure S7A). The divergent N-terminus was completely replaced, and that for 5 of the 6 differing amino acids the BLOSUM80 matrix gives a positive score for these substitutions ^120,121^. DNA and protein sequences of *RpS5b^RpS5a^* and *RpS5a^RpS5b^* in Table S1.

### Computational methods

The amino acid sequences of the 80 “canonical” ribosomal proteins in *D. melanogaster* were gathered from Flybase ^122,123^, and 12 *Drosophila* species covering over 70 million years of evolution were selected. One RP gene, *RpL41*, was excluded due to the difficulties posed by its short size (25 amino acids). For 10 of the species, we used genome assemblies that had been recently generated using HiC in conjunction with nanopore long-read sequencing, allowing almost complete assembly of full-length chromosome arms ^31^. For the remaining species, *D. mojavensis* and *D. grimshawi*, we used genome assemblies based on nanopore long-read sequencing only, as HiC-based assemblies were not available ^124,125^. Accession numbers and full details of genomes used are listed in Table S5. tBLASTn was carried out against the selected 12 Drosophila genomes, using the 78 *D. melanogaster* protein sequences as the input. The minimum protein sequence similarity threshold for BLAST hits was set at 25% and a maximum E-value was set at 3. A criterion was also included that any hit must cover at least 50% of the input protein sequence. This generated a list of hits for each of the 78 input genes in each of the 12 species (2155 hits in total). For each hit, the output contained information including the Muller element and coordinates of the hit, the coordinates of any gaps in the alignment, the overall protein sequence similarity to the input gene and the amino acids of the input gene to which the hit aligned, in addition to the E-value and protein sequence similarity score for each aligned segment of a hit. False positive hits that were not homologous to any ribosomal protein domain were manually removed, leaving 1373 hits. Comparison of hits from the same input gene in different species, using features such as the Muller element, sequence conservation and which part of the input gene the hit aligned to allowed us to deduce which hits corresponded to the same duplication event, and also identified which hits corresponded to the “canonical” gene in each species. Where multiple duplication events had led to multiple duplicates of a “canonical” gene being present in the same species, comparison of the sequences of each duplicate and the “canonical” gene was carried out to establish which was the “parent gene” for each duplication.

Next, we established which of the gaps in the alignments of hits represent the presence of introns. As the shortest intron that has been identified in D. melanogaster is 44bp ^126^, gaps shorter than 40bp were not classified as introns,. Gaps between 41bp and 200bp in length were checked manually for GT and AG dinucleotides at the start and end of each gap. Gaps between 201bp and 1000bp were classified as introns, whilst longer gaps were again checked manually for GT and AG dinucleotides. Comparison of the intron structure between duplicate and “parent gene” was used to classify the mode of duplication for each event. Where a “parent gene” had introns, duplication events which retained these introns in the same number and locations were classified as DNA-mediated. Duplications where the introns present in a “parent gene” were absent in the duplicate (even if new introns had been gained, in a handful of cases) were classified as retroduplications. In cases where the “parent gene” lacked introns, the mode of duplication could not be determined.

