## Supplementary Figures and Tables for "Ribosome abundance, not paralogue composition, is essential for germline development"

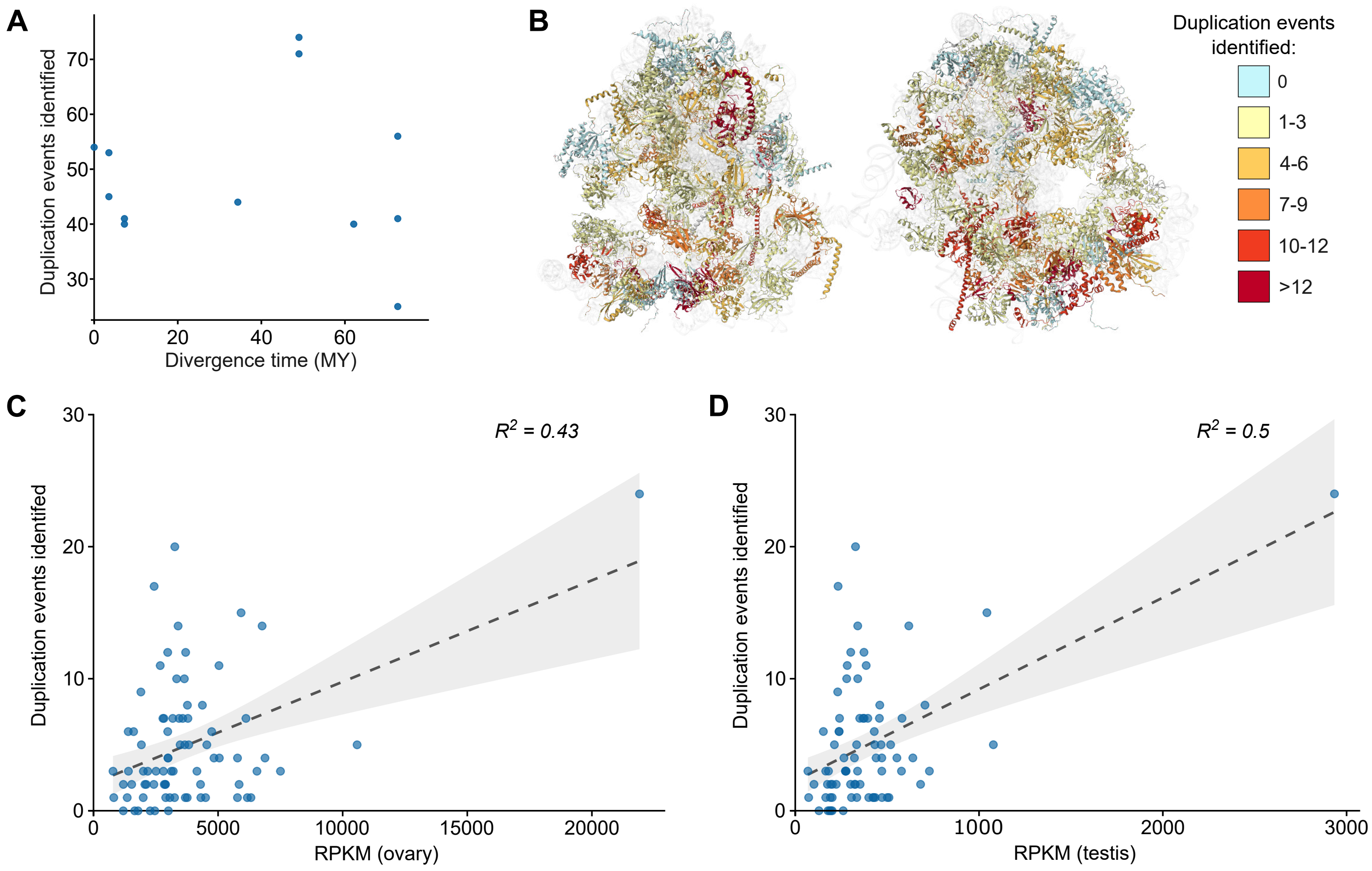


**Figure S1. Frequency of RP gene duplication correlates with germline expression (related to Figure 1)**

(**A**) Number of duplication events identified within each species, plotted against the divergence time of a given species from *D. melanogaster*. (**B**) Structure of the *D. melanogaster* ribosome (Anger *et al*., 2013; PDB: 4V6W) with RPs coloured by number of duplication events identified for each gene. rRNA is represented in light grey; . (**C-D**) Number of duplication events identified for each ribosomal protein gene plotted against its mRNA expression in the *D. melanogaster* adult (C) ovary or (D) testis. Expression data from Leader *et al*., 2018.

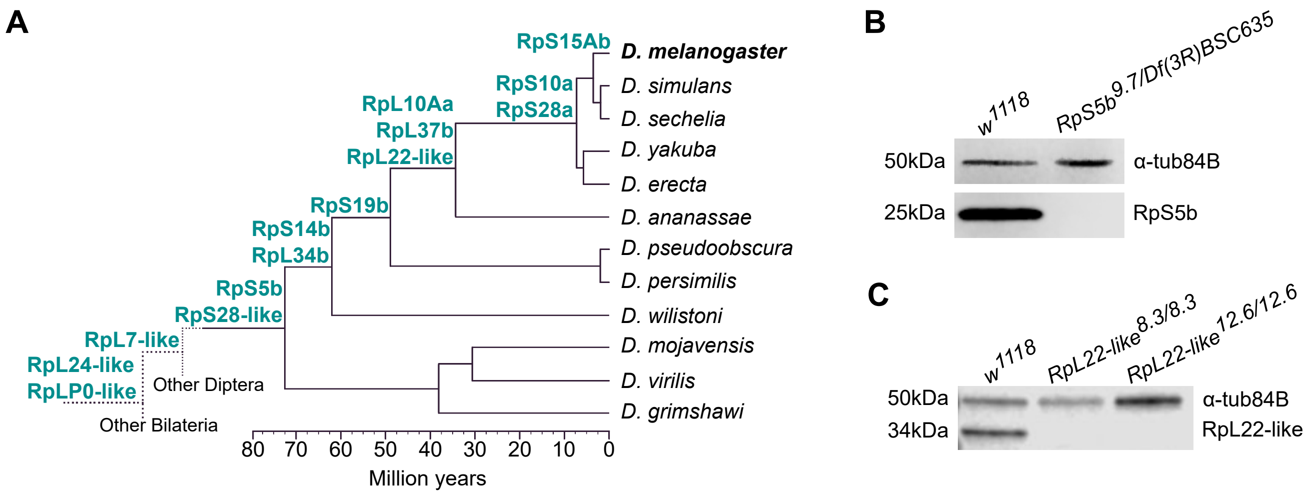


**Figure S2. RP paralogues with transcriptional activity in *D. melanogaster* (related to Figure 1)**

(**A**) The duplication event for each RP paralogue with transcriptional activity in *D. melanogaster* is depicted over the phylogeny of 12 Drosophila species. Phylogenetic tree scale based on Thomas and Hahn, 2017. (**B-C**) Western blot analyses with paralogue-specific antibodies against (B) RpS5b and (C) RpL22-like. (B) Comparison between $w^{1118}$ (control) and $RpS5b^{9.7/Df(3R)BSC635}$ ovaries, and (C) $w^{1118}$ (control), $RpL22\text{-}like^{8.3/8.3}$, and $RpL22\text{-}like^{12.6/12.6}$ testes.

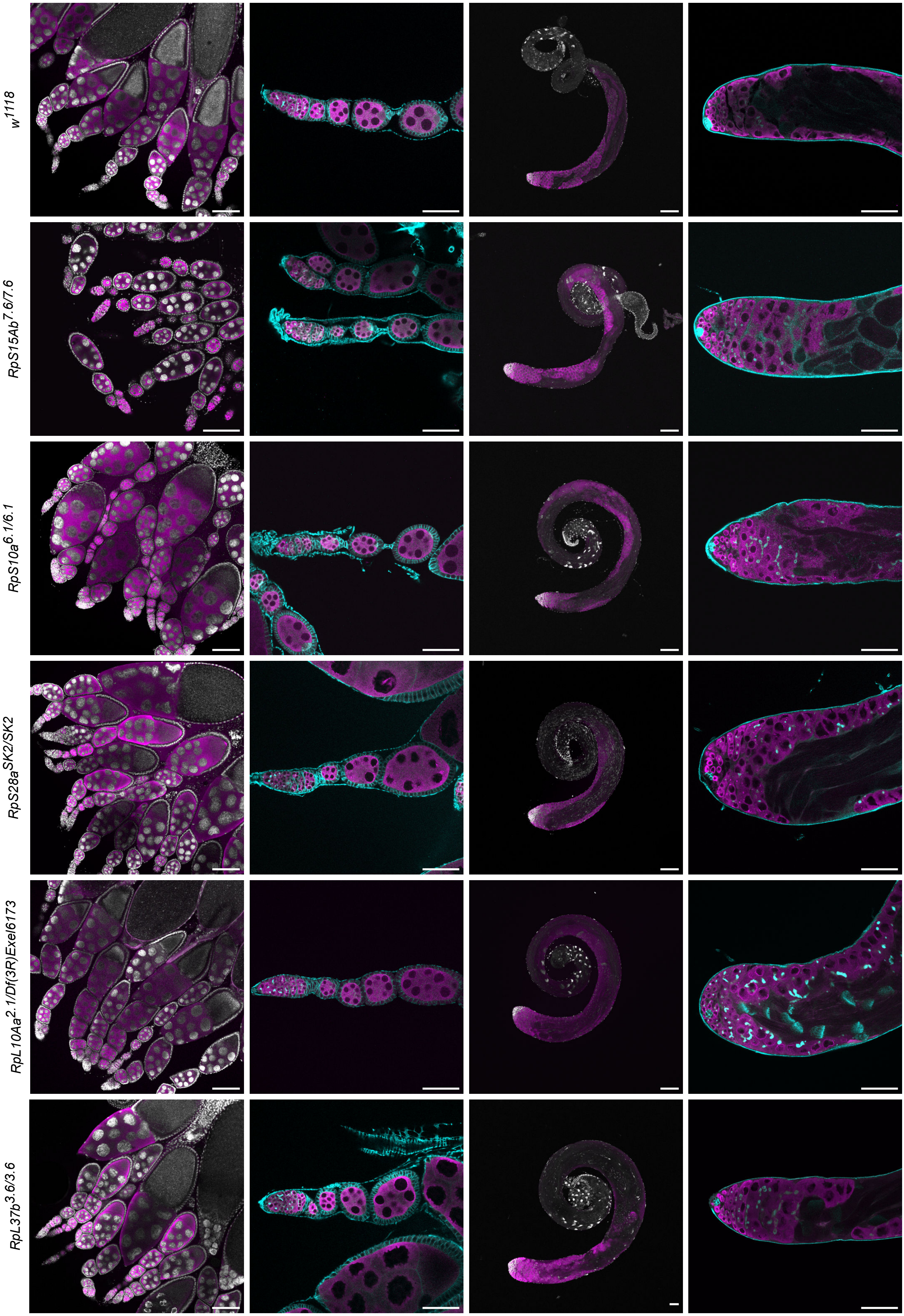

w1118
RpS15Ab7.6/7.6
RpS10a6.1/6.1
RpS28aSK2/SK2
RpL10Aa2.1/Df(3R)Exel6173
RpL37b3.6/3.6

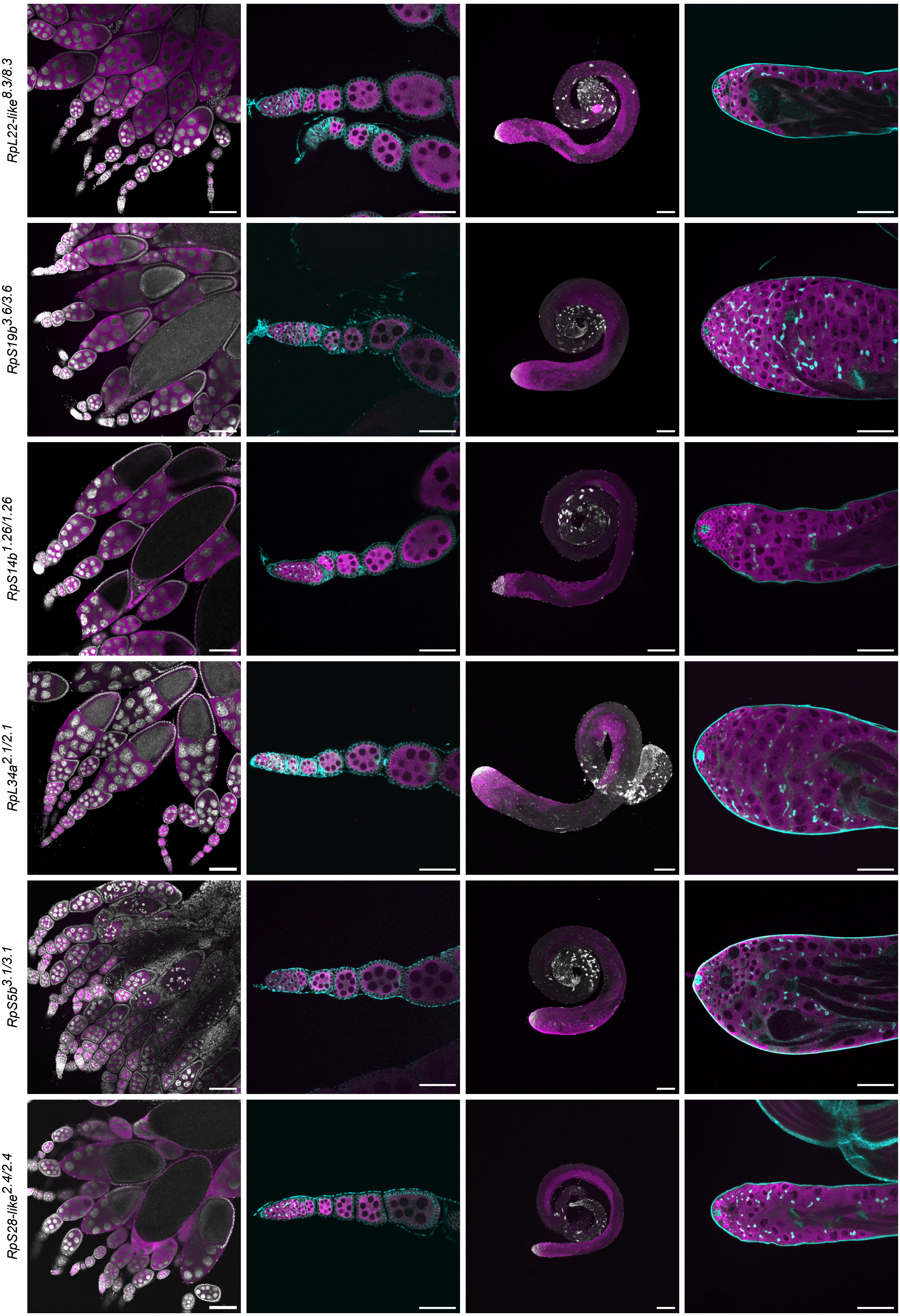
RpL22-like8.3/8.3
RpS19b3.6/3.6
RpS14b1.26/1.26
RpL34a2.1/2.1
RpS5b3.1/3.1
RpS28-like2.4/2.4

**Figure S3. The majority of “non-canonical” RP genes are dispensable for germline development (related to Figure 2)**

Representative ovaries (images in the left two columns) and testes (images in the right two columns) of $w^{1118}$ (control) and different “non-canonical” RP mutants labelled with DAPI (grey), vasa (germline, magenta) and α-spectrin (spectrosomes and fusomes, cyan). Testes were additionally labelled with fasciclin-III (hub cells, cyan). Scale bars = 100µm for images in the first and third columns, 50µm for images in the second and fourth columns.

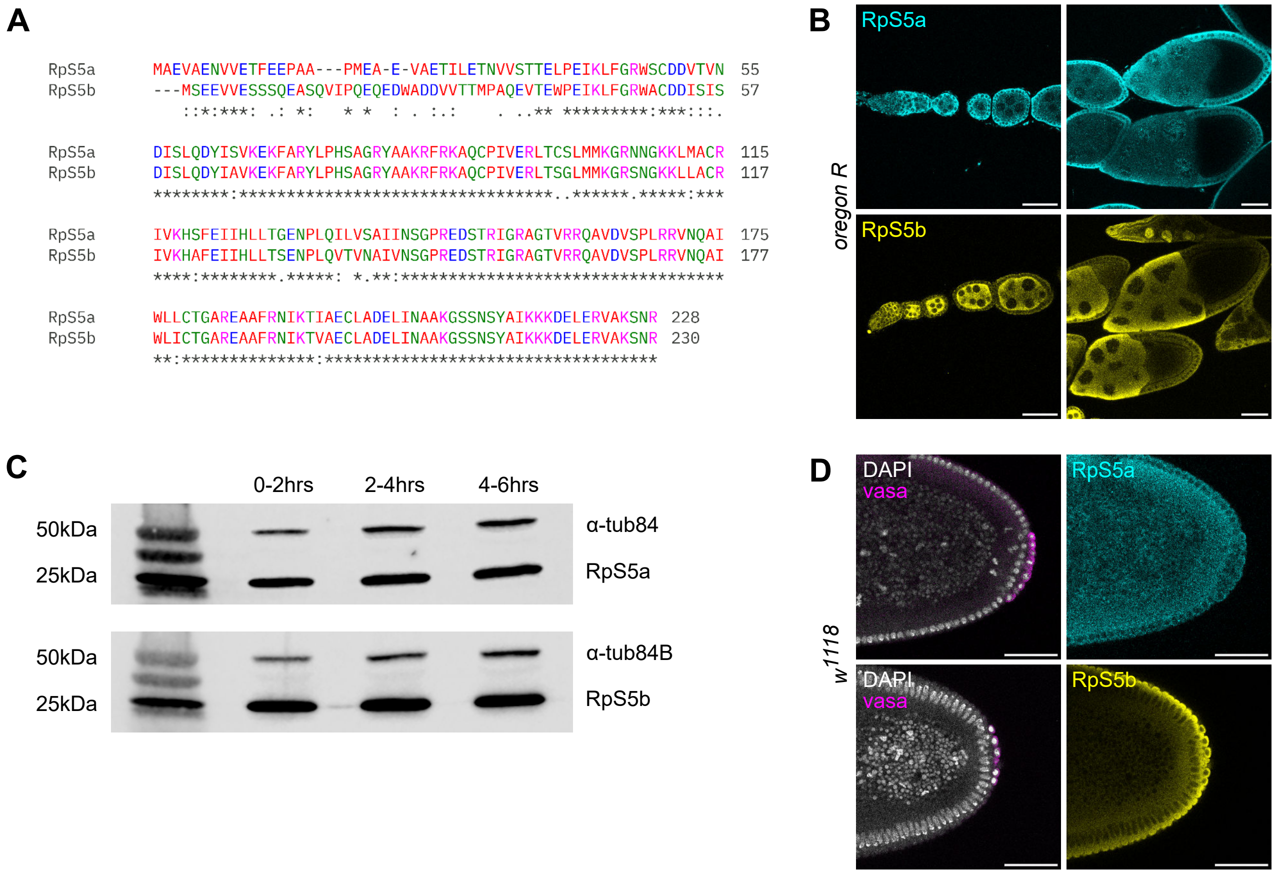


**Figure S4. *RpS5b* is predominantly expressed in germ cells and is maternally deposited into oocytes (related to Figure 2)**

(**A**) Alignment of *D. melanogaster* RpS5a and RpS5b protein sequences. (**B**) *Oregon R* ovaries (control), labelled with RpS5a (cyan) and RpS5b (yellow). Scale bars = 50µm. (**C**) Western blot analyses of $w^{1118}$ (control) embryos collected at subsequent 2-hour intervals post-fertilisation. (**D**) $w^{1118}$ blastoderm embryos (control), labelled with DAPI (grey), vasa (pole cells, magenta), RpS5a (cyan) and RpS5b (yellow). Scale bars = 50µm.

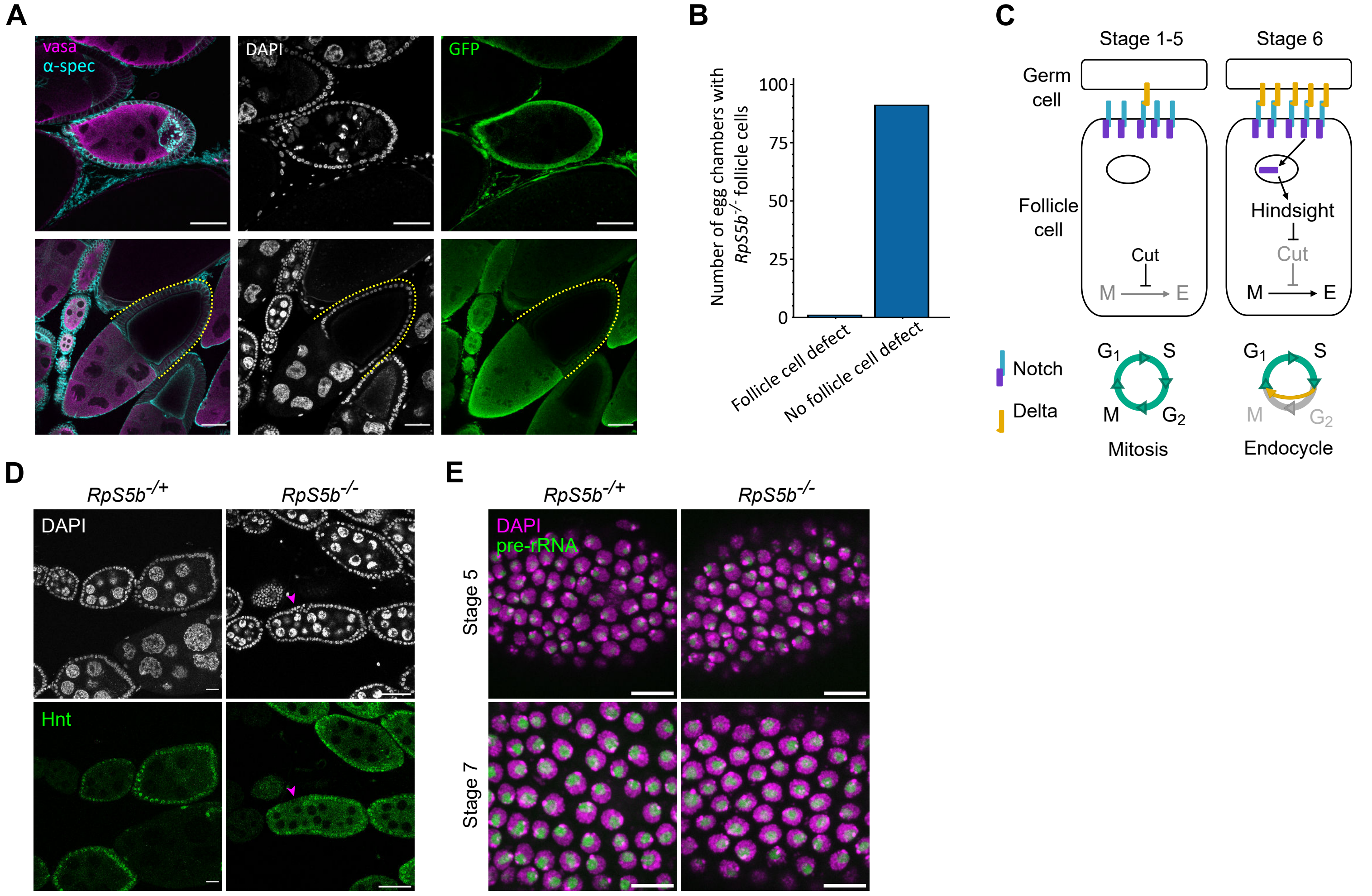


**Figure S5. Characterisation of cell type dependency using clonal analysis, and follicle cell mitotic-to-endocycle switch in *RpS5b$^{-/-}$* ovaries (related to Figure 3)**

(**A**) Representative confocal images of *hsFLP;; FRT82B RpS5b[3.1] / FRT82B Ubi-GFP* ovaries with examples of germline (top row) and somatic (bottom row) *RpS5b$^{-/-}$* clones, labelled with DAPI (DNA, grey), vasa (germline, magenta), α-spectrin (spectrosomes and fusomes, cyan) and GFP (green). Yellow dotted line indicates *RpS5b$^{-/-}$* follicle cells. Scale bars = 50µm. (**B**) Quantification of follicle cell multilayering phenotype in egg chambers containing somatic *RpS5b$^{-/-}$* clone. (**C**) Schematic showing Notch signaling before and after stage 6 of oogenesis. (**D**) Representative confocal images of *RpS5b$^{-/+}$* and *RpS5b$^{-/-}$* ovaries, labelled with DAPI (grey) and Hindsight (Hnt, green). Pink arrowheads indicate follicle cells in which Hnt signal is not observed at stage 6. Scale bars = 50µm. (**E**) Representative confocal maximum projections images (7-slice Z-stacks total, 1µm thick each) of follicle cells in egg chambers at stage 5 and stage 7 in *RpS5b*$^{-/+}$ and *RpS5b*$^{-/-}$ ovaries, labelled with DAPI (DNA, magenta) and pre-rRNA smFISH signal (nucleolus, green). Scale bars = 10µm.

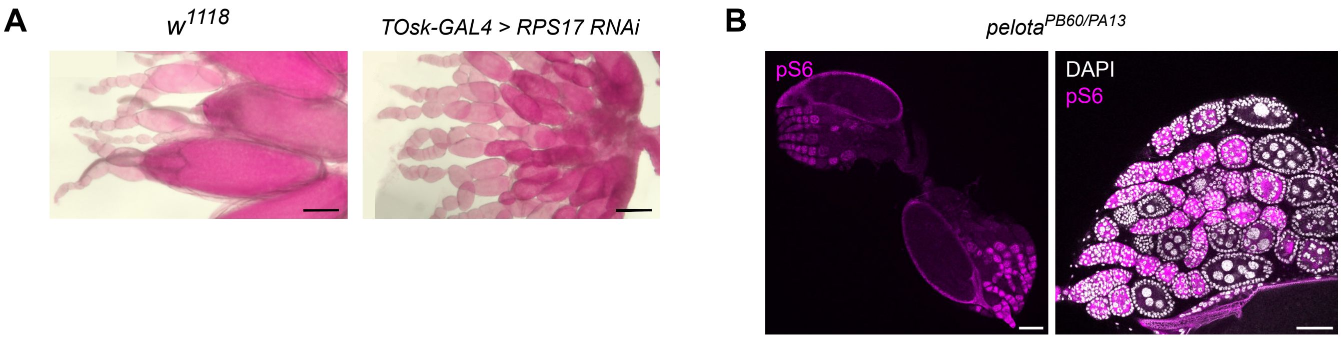


**Figure S6. Germline perturbation of "canonical" RP or ribosome recycling factor mimics *RpS5b*$^{-/-}$ phenotype (related to Figure 6)**

(**A**) Representative brightfield microscopy images of *w*$^{1118}$ (control) and *TOsk-GAL4 > RpS17 RNAi* ovaries stained with Periodic Acid Schiff's reagent. Pink staining shows glycogen accumulation. Scale bars = 100µm. (**B**) Representative confocal images of *pelota*$^{PB60/PA13}$ ovaries, labelled with DAPI (grey) and pS6 (magenta). Scale bars = 100µm for left image, 50µm for right image.

**A**

```
RpS5b-WT         MSEEVVESSSQEASQVIPQEQEDWADDVVTTMPAQEVTEWPEIKLFGRWACDDISISDIS 60
RpS5a[RpS5b]     MSEEVVESSSQEASQVIPQEQEDWADDVVTTMPAQEVTEWPEIKLFGRWACDDISISDIS 60
                 ************************************************************

RpS5b-WT         LQDYIAVKEKFARYLPHSAGRYAAKRFRKAQCPIVERLTSGLMMKGRSNGKKLLACRIVK 120
RpS5a[RpS5b]     LQDYIAVKEKFARYLPHSAGRYAAKRFRKAQCPIVERLTSGLMMKGRSNGKKLLACRIVK 120
                 ************************************************************

RpS5b-WT         HAFEIIHLLTSENPLQVTVNAIVNSGPREDSTRIGRAGTVRRQAVDVSPLRRVNQAIWLI 180
RpS5a[RpS5b]     HAFEIIHLLTSENPLQILVSAIINSGPREDSTRIGRAGTVRRQAVDVSPLRRVNQAIWLL 180
                 ****************: *.**:************************************:

RpS5b-WT         CTGAREAAFRNIKTVAECLADELINAAKGSSNSYAIKKKDELERVAKSNR 230
RpS5a[RpS5b]     CTGAREAAFRNIKTIAECLADELINAAKGSSNSYAIKKKDELERVAKSNR 230
                 **************:***********************************
```

**B**

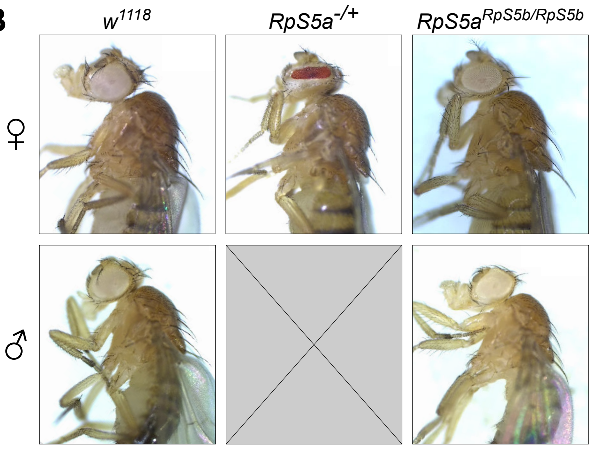


**Figure S7. RpS5a and RpS5b are interchangeable with regards to bristle development (related to Figure 7)**

(**A**) Alignment of wild type RpS5b protein and chimaeric RpS5 protein encoded by RpS5a$^{RpS5b}$. Purple box highlights N-terminal tail. (**B**) Comparison of females (top row images) and males (bottom row images) for $w^{1118}$, $RpS5a^{-/+}$ and $RpS5a^{RpS5b/RpS5b}$.

**Table S1. DNA and protein sequences of mutants generated in this paper**

tableS1.xlsx

**Table S2. RpS5b gene and protein sequences in Drosophila species analysed**

tableS2.xlsx

**Table S3. Probes used for pre-rRNA smFISH**

| Name | Sequence |
|---|---|
| prerRNA_mid_1 | ACCCATCTTCGTTTTATT |
| prerRNA_mid_2 | GCCCCACACTAACTATAT |
| prerRNA_mid_3 | TCTTTTTATGAGGTTGCC |
| prerRNA_mid_4 | CTCGCCACAAATACGCAC |
| prerRNA_mid_5 | AGGAAACGCCGTTGTTGT |
| prerRNA_mid_6 | TTCGACTTCCACTTTCGA |
| prerRNA_mid_7 | TCGCAACGCGTGTATAGT |
| prerRNA_mid_8 | CAGGTTGTTGCATTAGCC |
| prerRNA_mid_9 | AACCTGATGGATGCCAGG |
| prerRNA_mid_10 | TGGGTGACACATACTGCA |
| prerRNA_mid_11 | TCTGGTTGGTTATGGGGT |

**Table S4. Oligos used for CRISPR/Cas9**

| Gene | Sequence | |
|---|---|---|
| *RpS15Ab* | sense | CTTCGATCGTCGAGGATCACCGTGC |
| | antisense | AAACGCACGGTGATCCTCGACGATC |
| | sense | CTTCGTGACAACGATCTTGCCGGCA |
| | antisense | AAACTGCCGGCAAGATCGTTGTCAC |
| *RpS10a* | sense | CTTCGATGCAGTCGCTGAACTCACG |
| | antisense | AAACCGTGAGTTCAGCGACTGCATC |
| | sense | CTTCGATATTCGACCCGTCCGGTTC |
| | antisense | AAACGAACCGGACGGGTCGAATATC |
| *RpL10Aa* | sense | CTTCGCTCCAGGCAGTCTGGTCCTT |
| | antisense | AAACAAGGACCAGACTGCCTGGAGC |
| | sense | CTTCGAGGACCAGACTGCCTGGAGA |
| | antisense | AAACTCTCCAGGCAGTCTGGTCCTC |
| *RpL22-like* | sense | CTTCGCTACAGGAGTCACCTTAGCC |
| | antisense | AAACGGCTAAGGTGACTCCTGTAGC |

| | | |
|---|---|---|
| | sense | CTTCGTCAGCCTGCGTGATTGGTTG |
| | antisense | AAACCAACCAATCACGCAGGCTGAC |
| *RpL37b* | sense | CTTCGCCAAGGGTCGCAAGGCGCA |
| | antisense | AAACTGCGCCTTGCGACCCTTGGC |
| | sense | CTTCGAGGTTAGGAGCTTATAATC |
| | antisense | AAACGATTATAAGCTCCTAACCTC |
| *RpS19b* | sense | CTTCGTACAGGTGTCTCATAATAG |
| | antisense | AAACCTATTATGAGACACCTGTAC |
| | sense | CTTCGCTTCATATAGACGGCCTGTT |
| | antisense | AAACAACAGGCCGTCTATATGAAGC |
| *RpL34a* | sense | CTTCGTCGTCTCAGAGTCAAGCGT |
| | antisense | AAACACGCTTGACTCTGAGACGAC |
| | sense | CTTCGCATGCGCCTCTTGTTGGAGC |
| | antisense | AAACGCTCCAACAAGAGGCGCATGC |
| *RpS5b* | sense | CTTCGTTCTCCGGCTACCTCATCTT |
| | antisense | AAACAAGATGAGGTAGCCGGAGAAC |

| | | |
|---|---|---|
| | sense | CTTCGCAGCAGCCAGGAGGCGAGCC |
| | antisense | AAACGGCTCGCCTCCTGGCTGCTGC |
| | sense | CTTCGCGGCCCAGGAGGTCACCGAG |
| | antisense | AAACCTCGGTGACCTCCTGGGCCGC |
| | sense | CTTCGCTGCAGCGAGATGTCGCTGA |
| | antisense | AAACTCAGCGACATCTCGCTGCAGC |
| *RpS28-like* | sense | CTTCGCCTTGGAAGCGACAGGTTG |
| | antisense | AAACCAACCTGTCGCTTCCAAGGC |
| | sense | CTTCGAACTTCGGTTAGAATGCCT |
| | antisense | AAACAGGCATTCTAACCGAAGTTC |
| *RpS14b* | sense | CTTCGCTGCAGAATGGCTCCAAGAA |
| | antisense | AAACTTCTTGGAGCCATTCTGCAGC |
| | sense | CTTCGCGACACCTTCGTCCATGTCA |
| | antisense | AAACTGACATGGACGAAGGTGTCGC |

**Table S5. Genomes used for bioinformatic analysis**

| **Species** | **Strain** | **Accession** | **Reference** |
|---|---|---|---|
| *D. melanogaster* | iso-1 | PRJNA1119277 | Gebert *et al.*, 2024 |
| *D. simulans* | k-abb098 | PRJNA1119277 | Gebert *et al.*, 2024 |
| *D. sechellia* | k-s10 | PRJNA1119277 | Gebert *et al.*, 2024 |
| *D. yakuba* | k-s03 | PRJNA1119277 | Gebert *et al.*, 2024 |
| *D. erecta* | k-s02 | PRJNA1119277 | Gebert *et al.*, 2024 |
| *D. ananassae* | E-11001 | PRJNA1119277 | Gebert *et al.*, 2024 |
| *D. pseudoobscura* | k-s12 | PRJNA1119277 | Gebert *et al.*, 2024 |
| *D. persimillis* | k-s11 | PRJNA1119277 | Gebert *et al.*, 2024 |
| *D. willistoni* | k-s13 | PRJNA1119277 | Gebert *et al.*, 2024 |
| *D. mojavensis* | 15081-1352.22 | SRR6425997 | Miller *et al.*, 2018; Kim *et al.*, 2021 |
| *D. virilis* | k-s14 | PRJNA1119277 | Gebert *et al.*, 2024 |
| *D. grimshawi* | 15287-2541.00 (caf1) | SRR7642854,SRR7642855 | Kim *et al.*, 2021 |
