## Supplementary material for "Ribosome abundance, not paralogue composition, is essential for germline development": Key resource table

### KEY RESOURCES TABLE

| REAGENT or RESOURCE | SOURCE | IDENTIFIER |
|---|---|---|
| Antibodies | | |
| rabbit anti-vasa | Ruth Lehmann | |
| mouse anti-α-spectrin | DSHB | Cat#3A9; RRID:AB_528473 |
| mouse anti-fasciclin-III | DSHB | Cat#7G10; RRID:AB_528238 |
| mouse anti-hindsight | DSHB | Cat#1G9; RRID:AB_528278 |
| mouse anti-cut | DSHB | Cat#2B10; RRID:AB_528186 |
| rabbit anti-cleaved caspase-3 | Cell Signalling | Cat#9661; RRID:AB_2341188 |
| rabbit anti-phospho-H3 | Sigma-Aldrich | Cat#06-570; RRID:AB_310177 |
| rabbit anti-phospho-4E-BP | Cell Signalling | Cat#2855; RRID:AB_560835 |
| rabbit anti-phospho-RpS6 | Our lab | Gui *et al*., 2023 [20]; Romero-Pozuelo *et al*., 2017 [76] |
| rabbit anti-RpS5a | Paul Lasko | Kong *et al*., 2019 [43] |
| rabbit anti-RpS5b | Paul Lasko | Kong *et al*., 2019 [43] |
| rabbit anti-RpL22 | Julie Aspden | Hopes *et al*., 2022 [24] |
| rabbit anti-RpL22-like | Julie Aspden | Hopes *et al*., 2022 [24] |
| Goat anti-Rabbit IgG (H+L) Cross-Adsorbed Secondary Antibody, Alexa Fluor 568 | Invitrogen | Cat#A11011; RRID:AB_143157 |
| Goat anti-Mouse IgG (H+L) Highly Cross-Adsorbed Secondary Antibody, Alexa Fluor Plus 488 | Invitrogen | Cat#A32723; RRID:AB_2633275 |
| Donkey anti-Rat IgG (H+L) Highly Cross-Adsorbed Secondary Antibody, Alexa Fluor Plus 647 | Invitrogen | Cat#A48272; RRID:AB_2893138 |
| Goat anti-Chicken IgY (H+L) Secondary Antibody, Alexa Fluor™ 488 | Invitrogen | Cat#A11039; RRID:AB_2534096 |
| Alexa Fluor 647-AffiniPure Donkey Anti-Rabbit IgG (H+L) | Stratech | Cat#711-605-152; RRID:AB_2492288 |
| Goat anti-Mouse IgG (H+L) Highly Cross-Adsorbed Secondary Antibody, Alexa Fluor 647 | Invitrogen | Cat#A21236; RRID:AB_2535805 |
| Donkey anti-Rat IgG (H+L) Highly Cross-Adsorbed Secondary Antibody, Alexa Fluor 594 | Invitrogen | Cat#A21209; RRID:AB_2535795 |
| rabbit anti-α-tub84B | Abcam | Cat#ab15246; RRID:AB_301787 |
| IRDye 680LT Goat anti-Mouse IgG | LI-COR | Cat#926-68020; RRID:AB_10706161 |
| IRDye 800CW Goat anti-Rabbit IgG | LI-COR | Cat#926-32211; RRID:AB_621843 |
| Bacterial and virus strains | | |
| 5-alpha Competent E. coli (High Efficiency) | NEB | Cat#C2987H |

| | | |
|---|---|---|
| Biological samples | | |
| Chemicals, peptides, and recombinant proteins | | |
| Critical commercial assays | | |
| Periodic Acid-Schiff (PAS) Staining System | Sigma-Aldrich | Cat#395B-1KT |
| Deposited data | | |
| Experimental models: Organisms/strains | | |
| *D. melanogaster* : RpS28a$^{SK5}$ : *y[1] w[1118];; CG15527[SK5] / TM6c* | Eric Lai | Kondo *et al*., 2017 [36] |
| *D. melanogaster* : RpS28a$^{SK2}$ : *y[1] w[1118];; CG15527[SK2] / TM6c* | Eric Lai | Kondo *et al*., 2017 [36] |
| *D. melanogaster* : RpS5a$^{1}$ : *RpS5a[1] f[1]/FM6* | BDSC | #72 |
| *D. melanogaster* : w : *w[1118]* | Ruth Lehmann | |
| *D. melanogaster* : oregon R | University of Cambridge Fly Facility | |
| *D. melanogaster* : tj-GAL4 : *y[*] w[*]; P{GawB}NP1624 / CyO* | KDSC | #104055 |
| *D. melanogaster* : TOsk-GAL4 : *w; P{w[+mC]=matalpha4-GAL-VP16}V2H, P{w[+mC]=osk-GAL4::VP16}A11/CyO;* | VDRC | #314033 |
| *D. melanogaster* : Mat-tub-GAL4 : *w[*]; P{w[+mC]=matalpha4-GAL-VP16}V2H* | BDSC | #7062 |
| *D. melanogaster* : UAS-Dcr2 Nos-GAL4 : *UAS-Dcr2; nosp-ngt-gal4; nos-gal4::vp16* | Ruth Lehmann | Sanchez *et al*., 2016 [18] |
| *D. melanogaster* : RpS17 RNAi : *y[1] sc[*] v[1] sev[21]; P{y[+t7.7] v[+t1.8]=TRiP.HMS02492}attP2* | BDSC | #42656 |
| *D. melanogaster* : RpS18 RNAi : *; RpS18 RNAi* | VDRC | #108882 |
| *D. melanogaster* : RpL35 RNAi : *; RpL35 RNAi* | VDRC | #109425 |
| *D. melanogaster* : UTP6 RNAi : *; UTP6 RNAi* | VDRC | #34256 |
| *D. melanogaster* : tsc1 RNAi : *y[1] sc[*] v[1] sev[21]; P{y[+t7.7] v[+t1.8]=TRiP.GL00012}attP2* | BDSC | #35144 |
| *D. melanogaster* : mCherry RNAi : *y[1] sc[*] v[1];; P{VALIUM20-mCherry}attP2* | BDSC | #35785 |
| *D. melanogaster* : Udd$^{1}$ : *; Udd[1]/CyO; Dr/TM3* | Michael Buszczak | Zhang *et al*., 2014 [19] |
| *D. melanogaster* : Xrp1 : *w[*];; P{ry[+t7.2]=neoFRT}82B Xrp1[M2-73]* | BDSC | #81270 |
| *D. melanogaster* : RpS12$^{D97}$ : *w; RpS12[D97]* | Nicholas Baker | Ji *et al*., 2019 [61] |
| *D. melanogaster* : p53 : *y[1] w[1118]; p53[11-1B-1]* | BDSC | #6816 |
| *D. melanogaster* : FRT 82B : *;; P{neoFRT}82B* | Golnar Kolahgar | |
| *D. melanogaster* : FRT 82B GFP : *P{ry[+t7.2]=hsFLP}22, y[1] w[*];; P{neoFRT}82B P{Ubi-GFP.D}83* | Remi-Xavier Coux | Coux *et al*., 2018 [127] |
| *D. melanogaster* : pelota$^{PB60}$ : *w[*]; pelo[PB60] / CyO* | BDSC | 68149 |
| *D. melanogaster* : pelota$^{PA13}$ : *w[*]; pelo[PA13] / CyO* | BDSC | 68223 |

| | | |
|---|---|---|
| *D. melanogaster* : RpL22-like DF, RpL37b DF : *w; Df(2R) BSC769 / SM6a* | BDSC | 26866 |
| *D. melanogaster* : RpL37b DF : *w; Df(2R) Exel 7177 / CyO* | BDSC | 7906 |
| *D. melanogaster* : RpL34a DF : *w;; Df(3R) ED6232 / Tm6C Sb* | BDSC | 8105 |
| *D. melanogaster* : RpS10a DF : *w;; Df(3R) BSC499 / Tm6C Sb* | BDSC | 25003 |
| *D. melanogaster* : RpS5b DF :*w;; Df(3R) Exel 6172 / Tm6B Tb* | BDSC | 7651 |
| *D. melanogaster* : RpS5b DF : *w[1118]; Df(3R) BSC635/TM6C, Sb[1] cu[1]* | BDSC | 26505 |
| *D. melanogaster* : RpS19b DF : *w;; Df(3R) Exel 6196 / Tm6B Tb* | BDSC | 7675 |
| *D. melanogaster* : RpS15Ab DF : *w;; Df(3R) Exel 6196 / Tm6B Tb* | BDSC | 24352 |
| *D. melanogaster* : RpL10Aa DF : *w[1118]; Df(3R)Exel6173, P{w[+mC]=XP-U}Exel6173/TM6B, Tb[1]* | BDSC | 7652 |
| *D. melanogaster* : RpS28-like DF : *w[1118]; Df(2L)Exel8022 / CyO* | BDSC | 7813 |
| *D. melanogaster* : RpS28a DF : *w[1118]; Df(3R)BSC620 / TM6C, cu[1] Sb[1]* | BDSC | 25695 |
| *D. melanogaster* : RpL7-like DF : *w[1118]; Df(2L)BSC826/SM6a* | BDSC | 27900 |
| *D. melanogaster* : 2&3 balancer : *w; Sco / CyO; MKRS/TM6B* | Ruth Lehmann | |
| *D. melanogaster* : 2nd balancer : *w; Sco / CyO* | Ruth Lehmann | |
| *D. melanogaster* : 3rd balancer : *w;;TM2 / TM6C Sb* | University of Cambridge Fly Facility | |
| *D. melanogaster* : RpS15Ab$^{6.7}$ : *w; P{neoFRT}42D RpS15Ab[6.7] / CyO, Tb* | This paper | |
| *D. melanogaster* : RpS15Ab$^{7.7}$ : *w; P{neoFRT}42D RpS15Ab[7.7] / CyO, Tb* | This paper | |
| *D. melanogaster* : RpS10a$^{6.2}$ : *w;; P{neoFRT}82B RpS10a[6.2]/TM3 Sb* | This paper | |
| *D. melanogaster* : RpS10a$^{13.6}$ : *w;; P{neoFRT}82B RpS10a[13.6]/TM6B* | This paper | |
| *D. melanogaster* : RpL10Aa$^{2.1}$ : *w;; RpL10Aa[2.1] / TM6C Sb* | This paper | |
| *D. melanogaster* : RpL10Aa$^{5.7}$ : *w;; RpL10Aa[5.7] / TM6C Sb* | This paper | |
| *D. melanogaster* : RpL37b$^{3.1}$ : *w; P{neoFRT}42D RpL37b[3.1] / CyO, Tb* | This paper | |
| *D. melanogaster* : RpL37b$^{3.6}$ : *w; P{neoFRT}42D RpL37b[3.6] / CyO, Tb* | This paper | |
| *D. melanogaster* : RpL22-like$^{8.3}$ : *w; P{neoFRT}42D RpL22-like[8.3] / CyO* | This paper | |
| *D. melanogaster* : RpL22-like$^{12.6}$ : *w; P{neoFRT}42D RpL22-like[12.6] / CyO* | This paper | |
| *D. melanogaster* : RpS19b$^{1.5}$ : *w;; FRT82B 90Ew+ RpS19b[1.5] / TM6C Sb* | This paper | |
| *D. melanogaster* : RpS19b$^{3.3}$ : *w;; FRT82B 90Ew+ RpS19b[3.3] / TM6C Sb* | This paper | |

| | | |
|---|---|---|
| *D. melanogaster* : RpS14b$^{1.26}$ : *y[1] sc[1] v[1] RpS14b[1.26] / FM7i* | This paper | |
| *D. melanogaster* : RpL34a$^{5.8}$ : *w;; FRT82B 90Ew+ RpL34a[5.8] / TM6C Sb* | This paper | |
| *D. melanogaster* : RpL34a$^{8.5}$ : *w;; FRT82B 90Ew+ RpL34a[8.5] / TM6C Sb* | This paper | |
| *D. melanogaster* : RpS5b$^{3.1}$ : *w;; FRT82B 90Ew+ RpS5b[3.1] / TM6C Sb* | This paper | |
| *D. melanogaster* : RpS5b$^{6.4}$ : *w;; RpS5b[6.4] / TM6C Sb* | This paper | |
| *D. melanogaster* : RpS5b$^{8.2}$ : *w;; RpS5b[8.2] / TM6C Sb* | This paper | |
| *D. melanogaster* : RpS5b$^{9.7}$ : *w;; RpS5b[9.7] / TM6C Sb* | This paper | |
| *D. melanogaster* : RpS28-like$^{2.4}$ : *w; RpS28-like[2.4] / CyO* | This paper | |
| *D. melanogaster* : RpS28-like$^{3.1}$ : *w; RpS28-like[3.1] / CyO* | This paper | |
| *D. melanogaster* : RpS5b$^{8.2}$ Xrp1 : *w;; RpS5b[8.2] Xrp1[81270] / TM6C Sb* | This paper, combined with BDSC #81270 | |
| *D. melanogaster* : RpS5b$^{9.7}$ Xrp1 : *w;; RpS5b[9.7] Xrp1[81270] / TM6C Sb* | This paper, combined with BDSC #81270 | |
| *D. melanogaster* : RpS5b$^{9.7}$ RpS12$^{D97}$ : *w;; RpS5b[9.7] RpS12[D97] / TM6C Sb* | This paper, combined with RpS12$^{D97}$ from Nicholas Baker | |
| *D. melanogaster* : RpS5b$^{9.7}$ p53 : *w;; RpS5b[9.7] p53[6816] / TM6C Sb* | This paper, combined with BDSC #6816 | |
| *D. melanogaster* : RpS5b[RpS5a] : *w;; RpS5b[RpS5a] e / TM6C* | This paper | |
| *D. melanogaster* : RpS5a[RpS5b] : *y[1] sc[1] w v[1] RpS5a[RpS5b] / FM7h* | This paper | |
| Oligonucleotides | | |
| See table S2 for oligos used for CRISPR/Cas9. | This paper | |
| See table S3 for probes used for pre-rRNA smFISH. | This paper | |
| Recombinant DNA | | |
| pBluescript II Sk(+) | NovoPro | V011757 |
| Software and algorithms | | |
| Ribosomal protein BLAST pipeline | This paper | https://github.com/d-gebert/RP_blast |
| ImageJ | Schneider *et al.*, 2012 [128] | RRID:SCR_003070 |
| R Studio | | RRID:SCR_000432 |
| R package: ggplot2 | Wickham, 2016 [129] | RRID:SCR_014601 |
| R package: beeswarm | | RRID:SCR_024262 |
| R package: ggpubr | | RRID:SCR_021139 |
| Other | | |
